# Infection of Alfalfa Cotyledons by an Incompatible but Not a Compatible Species of *Colletotrichum* Induces Formation of Paramural Bodies and Secretion of EVs

**DOI:** 10.1101/2024.04.28.591504

**Authors:** Suchismita Ghosh, Kamesh C. Regmi, Barry Stein, Jun Chen, Richard J. O’Connell, Roger W. Innes

**Affiliations:** Department of Biology, Indiana University Bloomington, USA; Indiana University Bloomington Electron Microscopy Center, Indiana University Bloomington, USA; Université Paris-Saclay, INRAE, UR BIOGER, 91120 Palaiseau, France

**Keywords:** *Colletotrichum higginsianum*, *Colletotrichum destructivum*, *Medicago sativa*, Serial Block-Face Scanning Electron Microscopy (SBF-SEM), three-dimensional electron microscopy (3D-EM), Focused Ion Beam Scanning Electron Microscopy (FIB-SEM), IMOD, Multivesicular bodies (MVB), Paramural bodies, Extracellular Vesicles

## Abstract

Hemibiotrophic fungi in the genus *Colletotrichum* employ a biotrophic phase invading host epidermal cells followed by a necrotrophic phase spreading through neighboring mesophyll and epidermal cells. We used serial block face scanning electron microscopy (SBF-SEM) to compare subcellular changes that occur in *Medicago sativa* (alfalfa) cotyledons during infection by *Colletotrichum destructivum* (compatible on *M. sativa*) and *C. higginsianum* (incompatible on *M. sativa*). Three-dimensional reconstruction of serial images revealed that alfalfa epidermal cells infected with *C. destructivum* undergo massive cytological changes during the first 60 hours following inoculation to accommodate extensive intracellular hyphal growth. Conversely, inoculation with the incompatible species *C. higginsianum* resulted in no successful penetration events and frequent formation of papilla-like structures and cytoplasmic aggregates beneath attempted fungal penetration sites. Further analysis of the incompatible interaction using focused ion beam-scanning electron microcopy (FIB-SEM) revealed formation of large multivesicular body-like structures that appeared spherical and were not visible in compatible interactions. These structures often fused with the host plasma membrane, giving rise to paramural bodies that appeared to be releasing extracellular vesicles (EVs). Isolation of EVs from the apoplastic space of alfalfa leaves at 60h post inoculation showed significantly more vesicles secreted from alfalfa infected with incompatible fungus compared to compatible fungus, which in turn was more than produced by non-infected plants. Thus, the increased frequency of paramural bodies during incompatible interactions correlated with an increase in EV quantity in apoplastic wash fluids. Together, these results suggest that EVs and paramural bodies contribute to immunity during pathogen attack in alfalfa.

## Introduction

The *Colletotrichum* genus of fungal pathogens, which contains 280 accepted species, attacks over 3200 species of dicot and monocot species worldwide (Bhunjun et al., 2021; Crous et al., 2004; Liu et al., 2022; O’Connell et al., 2012; Yan et al., 2018). *Colletotrichum* infects many important food crops causing anthracnose leaf spot disease, which can cause large reductions in yield. Well-studied *Colletotrichum-*host interactions include *C. lindemuthianum* – bean (O’Connell et al., 1985), *C. graminicola* – maize (Mims & Vaillancourt, 2002; O’Connell et al., 2012), *C. higginsianum* – *Arabidopsis* (O’Connell et al., 2004), *C. sublineola* – sorghum (Wharton et al., 2001), *C. orbiculare*-cucumber (Xuei et al., 1988), and *C. destructivum* – alfalfa (Damm et al., 2014; Latunde-Dada et al., 1997). Of these, the infection of *Arabidopsis* by *C. higginsianum* is considered a model, because both plant and fungal partners can be genetically manipulated (O’Connell et al., 2004; Yan et al., 2018).

*Colletotrichum* begins its infection process by forming a melanized globular structure called the appressorium that arises from germ tubes produced by asexual fungal spores (conidia). Appressoria attach to leaf epidermal cells and form penetration pegs, allowing the fungus to push through epidermal cell walls, after which they form biotrophic hyphae that invaginate the host cell plasma membrane without rupturing it (Latunde-Dada et al., 1997; Mims & Vaillancourt, 2002; O’Connell, 1987; Wharton et al., 2001). This biotrophic phase is transient, lasting only 2-3 days on average (O’Connell et al., 2012; Yan et al., 2018). This is followed by a necrotrophic phase, where hyphae invade neighboring cells, killing the plant (Heath, 2000; Latunde-Dada et al., 1997). This biphasic mode of pathogenicity is called hemibiotrophy (Luttrell, 1974).

*C. higginsianum* expresses a large number of genes encoding secreted proteins during infection of *Arabidopsis*, including enzymes that mediate degradation of plant cell wall components such as cellulose, hemicellulose, and pectin (O’Connell et al., 2012). Expression of these enzymes is especially high during appressorium formation, indicating that these enzymes contribute to the penetration of the epidermal cell wall. During establishment of the biotrophic phase, genes encoding secondary metabolism enzymes and various effector proteins are induced (Kleemann et al., 2012). During the switch to necrotrophy, *C. higginsianum* starts to produce a large array of proteases, carbohydrate-active enzymes, and other lytic enzymes, as well as membrane transporters (O’Connell et al., 2012). During the necrotrophic phase, genes encoding necrosis-inducing proteins and many degradative enzymes such as proteases and cell wall-degrading enzymes are also up-regulated that cause tissue destruction and eventual cell death (Heath, 2000; Latunde-Dada et al., 1997; O’Connell et al., 2012).

Ultrastructural studies using electron microscopy have shown that *Colletotrichum* species often induce formation of electron-opaque structures called cell wall appositions or “papillae” beneath attempted sites of appresorial penetration. These papillae contain callose, a β-1,3-glucan polymer (Jacobs et al., 2003; Mims & Vaillancourt, 2002). Comparison of susceptible and resistant cultivars of French bean (*Phaseolus vulgaris*) infected with *C. lindemuthianum* showed that in both cases papillae formed beneath appressoria at similar frequencies, indicating that papillae formation is not specifically associated with unsuccessful penetration events (O’Connell et al., 1985). Similar studies between susceptible and resistant cultivars of *Sorghum bicolor* infected with *C. sublineolum* also revealed formation of papillae in both susceptible and resistant cultivars; however, the papillae formed in the resistant cultivar were more electron-opaque compared to the susceptible cultivar, suggesting deposition of additional materials in the resistant cultivar (Wharton et al., 2001). In both studies, infection of the susceptible cultivar began with the formation of an infection vesicle that arose from the appressorium and formed inside the host epidermal cell, giving rise to a primary biotrophic hypha. In the resistant cultivar, however, fewer infection vesicles were observed and formation of biotrophic hyphae was rare (O’Connell et al., 1985; Wharton et al., 2001).

The above studies focused on comparison of resistant and susceptible cultivars being infected by the same fungal strain. It is also informative to compare responses in a single host variety to infection by ‘compatible’ (adapted) and ‘incompatible’ (non-adapted) species of *Colletotrichum*. In this context, incompatible interactions are defined as interactions between a given fungal species that cannot infect any varieties of a given plant species (Heath, 2000). For example, *C. graminicola*, which normally infects maize, cannot infect any varieties of *Arabidopsis*, and is thus considered incompatible on *Arabidopsis*, whereas, *C. higginsianum* can infect most accessions of *Arabidopsis*, and is thus considered compatible on *Arabidopsis* (O’Connell et al., 2004; Shimada et al., 2006; Yan et al., 2018). Comparison of *Arabidopsis* infected with *C. higginsianum* to *Arabidopsis* infected with incompatible *Colletotrichum* species has shown that in the compatible interaction papillae were smaller and formed with lower frequency compared to the incompatible species (Shimada et al., 2006). Differences between compatible and incompatible responses were likewise reported in oats infected with *C. graminicola* (non-adapted) versus infection of maize with the same fungal strain (adapted) (Politis, 1976; Politis & Wheeler, 1973) suggesting that papillae contribute to immunity.

Formation of papillae is often associated with accumulation of multivesicular endosomes adjacent to the papillae (An, Huckelhoven, et al., 2006). These endosomes are typically ∼1 micron in diameter and contain multiple intraluminal vesicles of variable size. These endosomes are often referred to as multivesicular bodies (MVBs), but it should be noted that their appearance differs from that of MVBs associated with transport of plasma membrane receptors to lysosomes or vacuoles, which typically are less than 0.5 microns in diameter and contain smaller intraluminal vesicles with greater electron density and less variation in size (Buono et al., 2016). Multivesicular endosomes adjacent to papillae sometimes appear to fuse with the plasma membrane, at which point they are called paramural bodies (PMBs) (Marchants & Robards, 1968).

In this study, we used *Medicago sativa* (alfalfa) to compare subcellular responses to infection by compatible (*C. destructivum*) and incompatible (*C. higginisianum*) fungal species. These two species are phylogenetically very close, belonging to the same phylogenetic clade called the Destructivum species complex (Damm et al., 2014). All the members of this species complex share the same infection process, where the biotrophic hyphae are confined to one epidermal cell. In contrast, the biotrophic hyphae of most other hemibiotrophic *Colletotrichum* species extend into many host cells, including the mesophyll (da Silva et al., 2020; Damm et al., 2014). Examples include *C. lindemuthianum* and *C. orbiculare*, which belong to the Orbiculare species complex, and *C. sublineola* and *C. graminicola*, which belong to the Graminicola species complex (Bhunjun et al., 2021; Weir et al., 2012).

Alfalfa is commonly infected by *C. destructivum* (Damm et al., 2014; Latunde-Dada et al., 1997), which causes anthracnose disease in many forage and grain legume species including clover, alfalfa, cowpea, and lentil (Damm et al., 2014). *C. higginsianum,* which infects many crucifer species, including *Arabidopsis*, does not infect alfalfa. In this study, we wished to assess whether compatible and incompatible interactions in this system differed with regards to papilla formation and production of PMBs.

To assess papilla formation and PMBs, we performed three-dimensional electron microscopy using both serial block-face scanning electron microscopy (SBF-SEM) and focused ion beam scanning electron microscopy (FIB-SEM). All prior electron microscopy-based studies of plant-*Colletotrichum* interactions have used standard transmission electron microscopy (TEM), which provides two-dimensional cross-sections of infected cells (Kleemann et al., 2012; Mims & Vaillancourt, 2002; O’Connell et al., 1985; Wharton et al., 2001). The two-dimensional nature of these micrographs prevents complete understanding of the nature of infection structures observed. In contrast, SBF-SEM and FIB-SEM enable generation of 3D images by producing hundreds of serial sections in an automated manner, which can then be computationally assembled into a 3D structure (Denk & Horstmann, 2004; Xu et al., 2017). SBF-SEM and FIB-SEM differ from each other in the way that the serial sections are generated. The former uses a microtome housed inside the SEM to remove ∼40 nm sections from the sample block, with the fresh face imaged after removal of each section. FIB-SEM, in contrast, uses an ion beam to remove a 5-10 nm layer from the sample block between images. FIB-SEM imaging thus provides a higher resolution in the Z dimension. Both SBF-SEM and FIB-SEM reveal details such as the 3D shape and volume of the fungal hyphae, the topological nature of the circular objects observed in a single micrograph (e.g., spheres versus tubes), and the extent of membrane degradation throughout the volume of an infected plant cell, none of which are possible to assess from a single ultrathin section in classical TEM.

Using SBF-SEM and FIB-SEM, we were able to generate 3D models of the plant-fungal interface in both compatible and incompatible interactions. From these models, we obtained several important insights. At 24 hours post inoculation (hpi), both compatible and incompatible interactions displayed formation of fungal appressoria and induction of cytoplasmic vesicles and tubules around the periphery of the epidermal cell, with a higher concentration adjacent to appressoria. Neither interaction showed penetration of the plant cell wall by the fungus at this time point and ultrastructural differences between the two interactions appeared minor. At 60 hpi, in contrast, the differences were large. In the compatible interaction, biotrophic hyphae had largely displaced the central vacuole of epidermal cells, which requires a massive increase in host cell plasma membrane to accommodate these hyphae. While cytoplasmic vesicles and tubules were still abundant in the plant cell cytoplasm, no obvious papillae or PMBs were visible in the compatible interaction. In the incompatible interaction, no biotrophic hyphae or penetration pegs had formed, despite abundant appressoria being present. Notably, large cytoplasmic aggregates had formed underneath many appressoria, which we think are precursors of papillae, and numerous PMBs were observed spread around the periphery of epidermal cells. Consistent with the increase in PMBs, we also detected an increase in extracellular vesicle number in apoplastic wash fluids. Together, these observations support the assumption that papillae and extracellular vesicles contribute to non-host resistance in Medicago.

## RESULTS

### Both compatible and incompatible fungal species induce vesicle formation in host epidermal cells prior to penetration

We wanted to understand whether the infection of alfalfa with its compatible *Colletotrichum* species induces visible changes in host cell ultrastructure prior to the formation of biotrophic hyphae. We therefore chose 24 hpi to visualize these subcellular changes, as we never observed biotrophic hyphae at this time point.

To image subcellular structures, we used SBF-SEM and chemical fixation of samples. We employed chemical fixation rather than high pressure freezing in order to preserve appressoria. In prior SBF-SEM analyses of Arabidopsis cotyledons infected with *Colletotrichum higginsianum* (a compatible interaction), we observed that appressoria were entirely missing, despite the presence of abundant intracellular hyphae (Regmi et al., 2023). In this study, we wished to capture subcellular defense responses in incompatible interactions that lacked intracellular hyphae, and thus needed to be able to locate sites of attempted penetration. This proved impossible with high pressure freezing, hence our decision to employ chemical fixation, instead, despite the advantages of high pressure freezing for preserving membrane ultrastructure. A limitation of chemical fixation is that dehydration of cells during the fixation process can lead to changes in membrane structures. In our analyses below, we therefore focused on membrane structures that were absent in our non-infected controls, and when comparing compatible and incompatible interactions, structures that were unique to one or the other.

For the mock inoculation, 200 serial images were obtained (Fig. 1A and Supplementary Video S1; see end of Methods for link to Supplementary Videos) and for the compatible infection (*C. destructivum*), 201 serial images were obtained (Fig. 1B-1F and Supplementary Video S2). For both samples, this represents ∼8 µm of cell depth. The mock infected alfalfa cotyledon (Fig. 1A) showed an intact plant cell membrane as well as an intact tonoplast with very little intracellular vesiculation (enlarged in Fig. 1A1). In contrast, the *C. destructivum-*infected cotyledon displayed abundant intracellular vesiculation (Fig. 1B). As expected, no appressoria were present in the mock-infected sample (Fig. 1A), whereas dome-shaped appressoria were concentrated over anticlinal cell walls, in the valleys between epidermal cells of the infected cotyledon (Fig. 1B). This is expected because spores tend to accumulate in this location, and hence appressoria form there. Electron-opaque vesicular structures were visible that appeared to be near the vicinity of the plant cell plasma membrane. These structures could be grouped into two classes based on their level of staining: heavily stained (electron-opaque; yellow arrowheads) and lightly stained (blue arrowheads) (Fig. 1B and 1C). These electron-opaque circular structures were absent from mock-inoculated plant cells as seen in Fig. 1A and Supplementary <u>Video S1</u>.

**Fig. 1.**
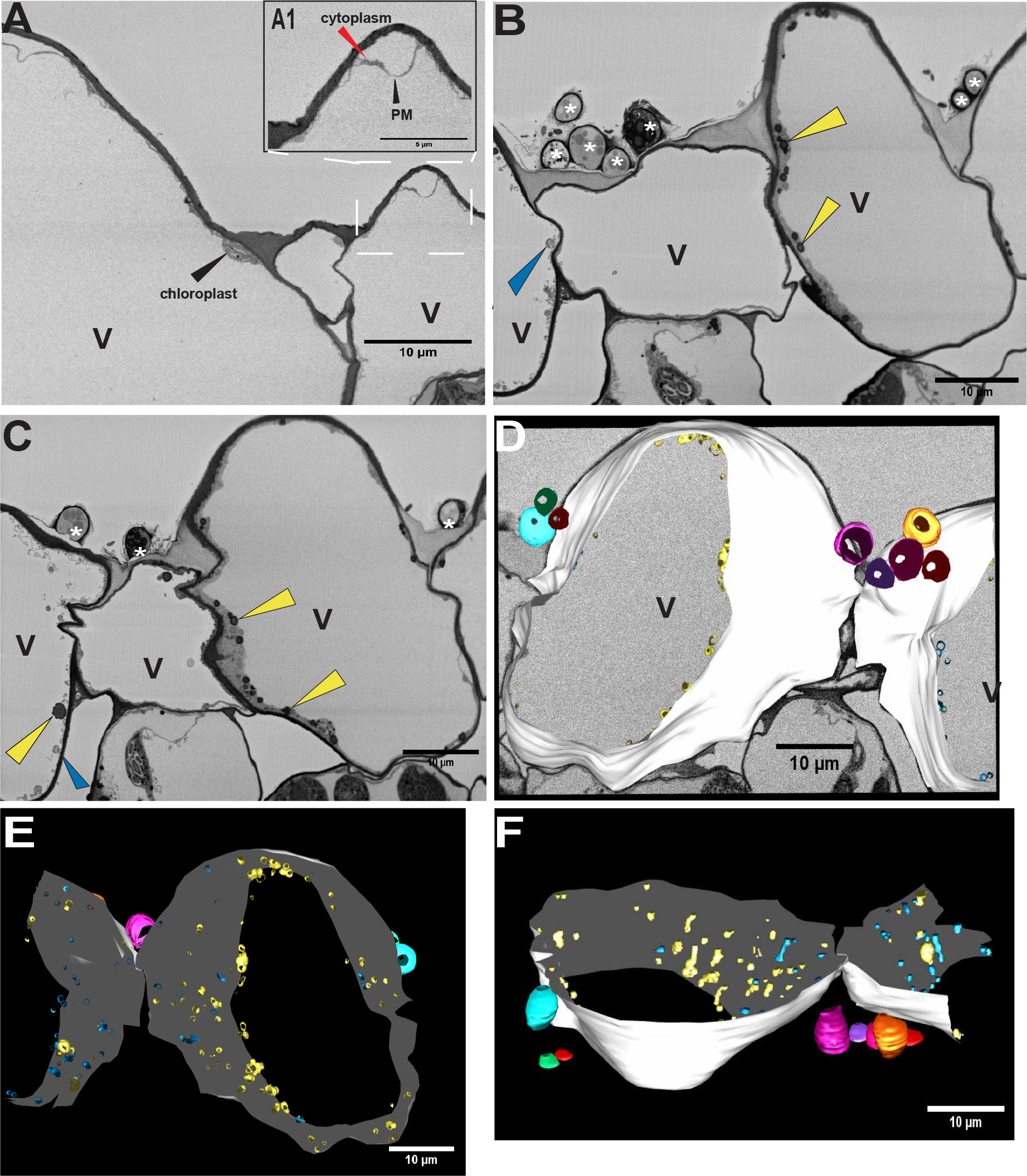
A compatible fungal pathogen induces vesiculation in host cells prior to formation of biotrophic hyphae. **A**, Mock-inoculated epidermal cells show very little vesiculation. Image shows the 1^st^ micrograph from a stack of 200 SBF-SEM images of three epidermal cells on an alfalfa cotyledon mock inoculated with water and imaged 60 hours after inoculation. Supplementary Video S1 shows all 200-images in this stack. Note large central vacuoles (V), and chloroplast visible in the epidermal cell. The boxed region is enlarged in **A1**, which shows the plasma membrane (PM) and thin peripheral cytoplasm. **B** and **C**, *C. destructivum* induces vesiculation by 24 hpi. Panels B and C show the 14^th^ and 84^th^ micrograph from an SBF-SEM image stack of alfalfa epidermal cells infected with *C. destructivum* (a compatible species) and imaged 24 hpi. Fungal appressoria are marked with an asterix and appear as dome shaped structures on the adaxial surface of host epidermal cells. Heavily-stained circular structures are indicated by yellow arrows. These structures appear mostly along the plasma membrane of the plant cell and are absent from mock-infected cotyledons shown in panel A. Blue arrows indicate moderately-stained circular structures. Supplementary Video S2 shows all 201-images in this stack. **D**, 3D model of two adjacent epidermal cells superimposed on a greyscale SBF-SEM image from the same stack shown in B and C. This model was generated using IMOD software. The plant cell wall is shown in white for the two adjacent cells. Fungal appressoria are shown as colored donut-shaped structures sitting above the plant cell. Heavily-stained vesiculo-tubular structures observed in images B and C are modelled in yellow, while moderately-stained structures are modelled in blue. **E**, 3D model of image stack., **F**, the same model rotated 90 degrees vertically. Please refer to Supplementary Video S3 to see this model rotating in space. V, vacuole; *, appressoria.

To assess the three-dimensional shape of the vesicle-like structures, we used the free software program IMOD (Kremer et al., 1996). Figure 1D shows a representative gray-scale image with the model superimposed on top of it, which is 180 degrees rotated from Figures 1B and 1C. The entire model is shown rotating in space in Supplementary Video S3. 3D imaging is required to distinguish between spherical and tubular structures that both appear as circles in two-dimensional sections. We modelled the circular structures shown in Figures 1B and 1C, maintaining the same color code. Figure 1E is a snapshot of the model which reveals that even though these structures appeared vesicular in a two-dimensional SEM image, some of them are in fact tubular rather than spherical and in this snapshot these tubular structures are roughly in the same orientation. Figure 1F shows the model rotated to better visualize their tubular nature. The electron-opaque structures (yellow) were more abundant than the less densely stained structures (blue). Both classes of structures appeared near the plasma membrane, suggesting that they may be a response to the fungal appressoria. Some of these structures were vesicular, as seen by their spherical shape, while some taper into tubes along the periphery of the plant cell under attack. Supplementary Figure S1B shows a single frame from an SBF-SEM image stack of a second compatible infection at 24 hpi, while Supplementary Video S4 shows the entire set of images. These images show the same pattern of vesiculation as observed in Supplementary Video S2.

Next, we examined the response of alfalfa to inoculation with an incompatible fungal species (*C. higginsianum*) to assess whether the response differs from a compatible strain at 24 h. We collected 251 serial images representing ∼10 µm of cell depth (Fig. 2 and Supplementary Video S5). Figure 2A shows the 111^th^ image of the image stack, while Figure 2B shows the 212^th^ image. Like its compatible counterpart (Fig. 1) we observed both electron-opaque and less dense vesicular structures (Fig. 2A and 2B, yellow and blue arrows). This indicates that the intracellular vesiculation observed in the compatible interaction also occurs during an incompatible interaction. In addition to the electron-opaque and less-dense vesicular structures, we observed a third type of circular structure that was more prominent in the incompatible interaction, which appeared to be membrane-bound circles with no visible internal staining (indicated by pink arrows in Fig. 2A). These structures were rarely observed in compatible interactions.

**Fig. 2.**
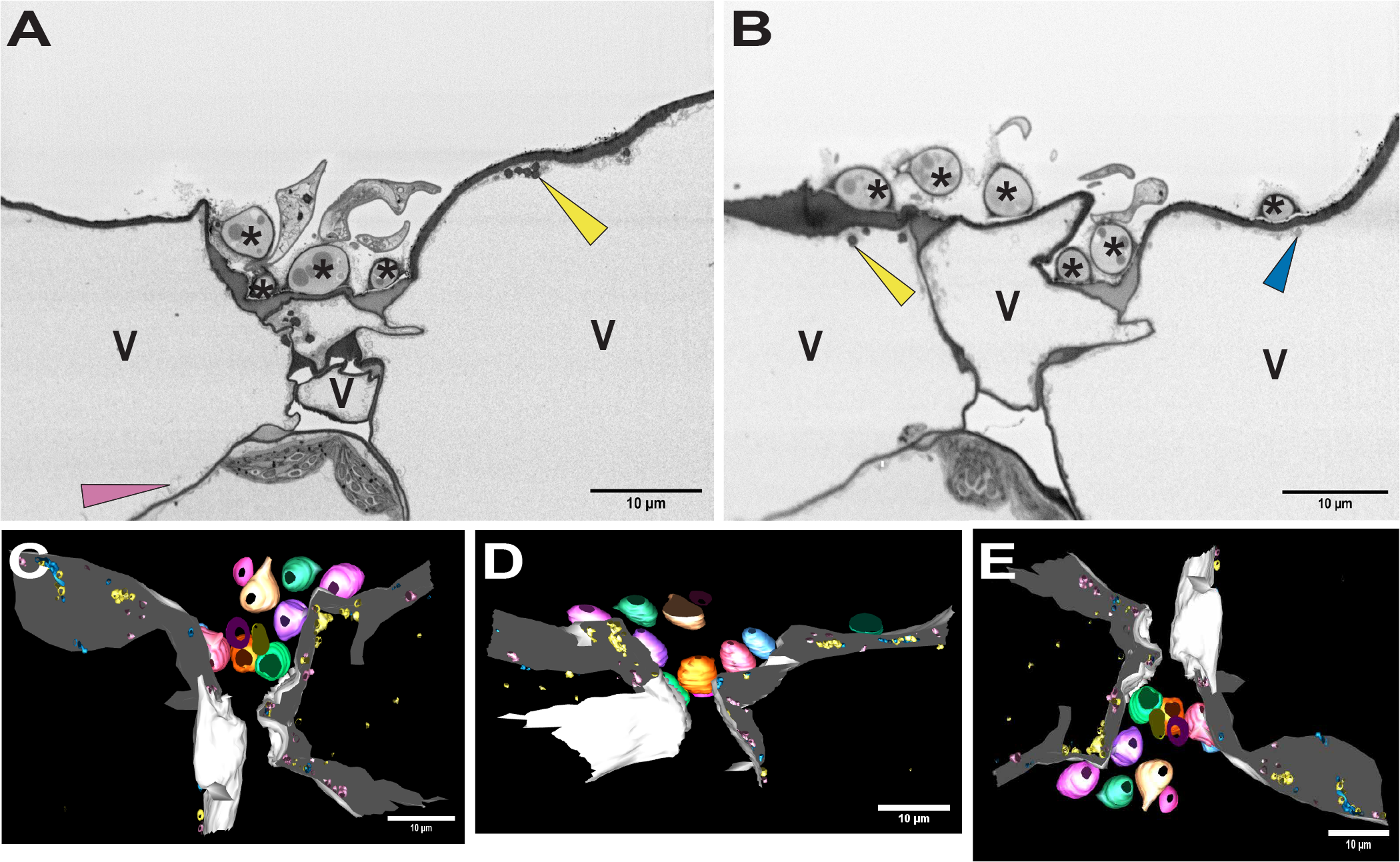
An incompatible fungal pathogen induces vesiculation in host cells, including a third class of vesicles not observed during a compatible interaction. **A** and **B**, the 111^th^ and 212^th^ micrographs from an SBF-SEM image stack containing 251 images showing alfalfa epidermal cells infected with *C. higginsianum* (incompatible on alfalfa) and imaged at 24 hpi. Labels are the same as described for Figure 1. Very lightly stained circular structures, which were not observed in the compatible interaction, are seen throughout the entire stack of 251 images and are marked with pink arrowheads in panel A. Heavily-stained circular structures are indicated by yellow arrowheads, while moderately-stained structures are indicated by blue arrowheads. Supplementary Video S5 shows all 251-images in this stack. **C**, 3D model of the entire volume of 251 SBF-SEM images. Supplementary Video S6 shows this 3D model rotating in space. **D**, and **E**, shows the same model in C rotated 90 degrees and 180 degrees vertically to better visualize the vesiculo-tubular structures. Heavily-stained structures are modeled in yellow, moderately- stained structures are modelled in blue, while lighter-stained structures are modelled in pink. V, vacuole; *, appressoria.

Figure 2C shows a snapshot of a 3D model generated from the entire image stack of the incompatible interaction. Supplementary Video S6 shows the full model. Figures 2D and 2E are snapshots of the model rotated vertically at 90 degrees and 180 degrees, respectively, to reveal the nature of the electron-opaque and less dense structures. Similar to the compatible interaction at 24 hours, penetration of plant cells by fungal hyphae was not observed, although numerous appressoria were present on the cell surface. Some of the electron-opaque structures formed a meshwork of tubules similar to that observed in the compatible interaction (yellow in the model). The lighter stained structures (blue) appeared to be more vesicular. The hollow-looking pink structures appeared to form both spherical and tubular structures. Supplementary Figure S1C and Supplementary Video S7 show a second incompatible inoculation of alfalfa imaged at 24 hours, revealing a similar pattern of vesiculation as observed in Supplementary Video S5.

Supplementary Figure S2A quantifies the number of vesicles and tubules observed in either compatible or incompatible interactions at 24 h for both the electron opaque (yellow) and the lightly stained (blue) structures across 20 µm of plant cell depth. Since the very lightly stained hollow structures (pink) were rarely visible or were too small to model in the compatible interaction, we did not quantify those particles. The 3D model was used to distinguish between the vesicles and the tubules. For our calculations, any structure that was twice as long as it was wide was considered a tubule. Both the compatible interaction (Supplementary Videos S2 and S4) and incompatible interaction (Supplementary Videos S6 and S7) were modelled and quantified for the graph. These analyses did not show any significant differences between the number of vesicles and tubules observed between the compatible and incompatible interactions. The electron opaque particles (yellow) seemed to form more tubules than vesicles, whereas the lightly stained particles (blue) formed more vesicles than tubules in both interaction types at 24h.

### *C. destructivum* biotrophic hyphae displace the central vacuoles of epidermal cells by 60 hpi

The above analyses revealed cellular responses occurring prior to formation of biotrophic hyphae but did not reveal large differences between compatible and incompatible interactions. We therefore imaged infection sites at 60 hpi. At this time-point, we expected to see extensive biotrophic hyphae in the compatible interaction, but very little cell death. In incompatible interactions, in contrast, we expected to see a lack of biotrophic hyphae, and possibly death of epidermal cells and/or formation of defensive structures such as papillae (cell wall appositions).

As seen in Figure 3, by 60 hpi, *C. destructivum* had extensively colonized the epidermal cell with its biotrophic hyphae, which largely displaced the central vacuoles of infected epidermal cells, although the plasma membrane and tonoplast remained intact. We were able to observe a penetration peg (black arrow in Fig. 3A), which is the site where the fungus penetrates the plant cell wall beneath the appressorium. This event could be captured because we imaged ∼16 µm of the plant cell volume (398 sections) using the SBF-SEM (Supplementary Video S8), enabling us to find the rare section that captured such an event. We used IMOD to reconstruct the imaging volume in 3D, which allowed us to visualize the 3D branching structures of individual hyphae. We could not visualize any interfacial matrix at this resolution (material surrounding biotrophic hyphae that is expected to accumulate between the plasma membranes of the fungus and the plant). The biotrophic hyphae were highly stained. Intense fragmentation of the plant vacuolar membrane was seen in the vicinity of the biotrophic hyphae, as well as near the surface of the plant cell (Fig. 3A1). In contrast, the plant chloroplasts, mitochondria, and plasma membrane remained intact, indicating that the fungus was still in its biotrophic phase. Figure 3B represents the 300^th^ micrograph from the same image stack, which shows the biotrophic hyphae almost displacing the entire volume of the plant vacuole, with the vacuole becoming highly fragmented (Fig. 3B1). Like the 24-hour time point, densely stained structures were also observed (indicated with yellow arrows in Figure 3B1 and modelled using the same color code). A 3D model of the entire imaging volume was first created without the vacuolar membrane to simplify the model (Supplementary Video S9). Next, we modelled the area of vacuolar membrane disintegration as mentioned above, which can be seen in Supplementary Video S10. Figure 3C shows a snapshot of the model with the biotrophic hyphae and the heavily stained yellow structures and the hollow structures in pink. For ease of visualization, biotrophic hyphae 3 and 4 are dotted to better see the yellow and pink structures along the periphery of the membrane. Figure 3D represents a gray-scale image superimposed on the model, which enabled us to connect the biotrophic hyphae as continuous structures occupying nearly the entire plant cell volume. Figure 3E is a 90-degree rotated image of the same model shown in Figure 3D. To capture the amount of membrane disintegration, the same model is rotated and shown in Figure 3F, showing the extensive nature of these structures, roughly in the same orientation much like Figure 1F. The heavier stained yellow structures appear more tubular and seem to form an interconnected mesh near the plant cell wall. The lighter stained structures are occasionally vesicular spheres, but mostly tubular; however, they seem to be less interconnected compared to the yellow ones. Overall, the fungus occupied the majority of the plant cell, suggesting massive expansion of the plant plasma and tonoplast membranes must occur to accommodate the very large biotrophic hyphae. Supplementary Figure S1D shows a second compatible infection of alfalfa imaged at 60 hpi, while Supplementary Video S11 shows the entire SBF-SEM image stack. These images confirm the extensive remodeling of the host cell vacuolar and plasma membranes during the compatible interaction.

**Fig. 3.**
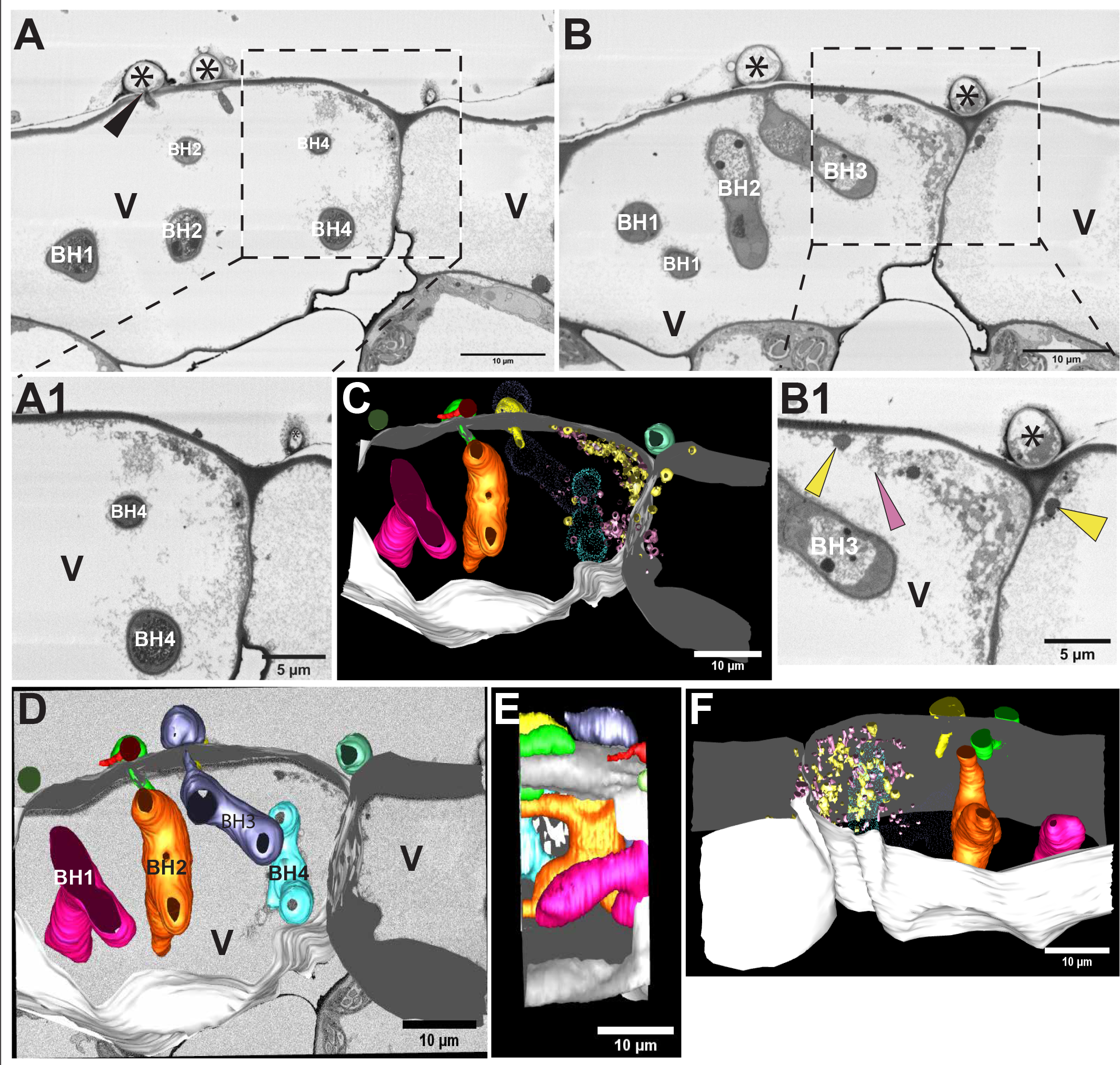
Alfalfa cotyledons accommodate extensive biotrophic hyphae at 60 hpi with *C. destructivum*. **A** and **B**, 100^th^ and 300^th^ micrograph, respectively, from an SBF-SEM image stack containing 398 images. Boxed regions in panels A and B are enlarged in panels A1 and B1, respectively. The black arrowhead in panel A marks a fungal penetration peg. Biotrophic hyphae (BH) are numbered and correspond to the biotrophic hyphae modeled in panel D. Heavily-stained circular structures are indicated with yellow arrowheads in panel B1. These structures appear mostly along the plasma membrane of the plant cell. The pink arrowhead in B1 indicates lighter stained circular structures. Supplementary Video S8 shows all 398 images in this stack. **C**, 3D model generated from this stack using IMOD. The plant cell wall is shown in white for the two adjacent cells. Note the extensive biotrophic hyphae and the vesiculo-tubular structures. The dark structures are modelled with yellow, while the lighter ones are modelled in pink. For the ease of visualization, BH3 and BH4 are dotted to better see the vesiculo-tubular structures near the plant plasma membrane. **D**, 3D model superimposed on a greyscale SBF-SEM image from the same stack. The biotrophic hyphae are numbered 1 to 4 and correspond to those labelled in panels A and B. Supplementary Video S9 shows this model rotating in space. **E**, 90-degree rotation of the 3D model showing the extensive hyphae extending across the depth of the epidermal cell. **F**, the same model shown in C rotated vertically to better visualize the vesiculo-tubular structures. Supplementary Video S10 shows a 3D model of these structures. BH, biotrophic hyphae; V, vacuole; *, appressoria.

Given the extensive membrane remodeling and vesiculation observed by SBF-SEM, we wished to determine whether cell death was occurring during the compatible interaction. We thus stained cotyledons with trypan blue (Fernández-Bautista et al., 2016; O’Connell et al., 2004) at both 24 and 60 hpi. We observed no evidence of cell death at either time point with the compatible strain *C. destructivum* (Supplementary Fig. S3), even though this interaction is expected to switch to a necrotic phase by 72 hpi.

### The incompatible fungus *C. higginsianum* fails to penetrate alfalfa epidermal cells

Attempted penetration of plant epidermal cells by non-adapted fungal species often induces formation of localized defense structures known as cell wall appositions or papillae (An, Ehlers, et al., 2006; An, Huckelhoven, et al., 2006; Rubiato et al., 2022). Extensive studies of barley infected with incompatible species of powdery mildew fungi have revealed formation of papillae or encasements as pre- and post-invasive defense structures respectively (Assaad et al., 2004; Bohlenius et al., 2010; Heitefuss & Ebrahim-Nesbat, 1986; Zeyen & Bushnel, 1979). The incompatible interaction interface in the hemibiotrophic fungus *Colletotrichum* is less studied. We imaged incompatible interactions at the 60-hour time point to compare the differences in host cell responses between compatible and incompatible *Colletotrichum* species.

Contrary to *C. destructivum* (compatible) (Fig. 3), *C. higginsianum* (incompatible) was unable to penetrate alfalfa epidermal cells even at 60 hpi (Fig. 4). Interestingly, as seen in Figure 4A and Supplementary Video S12, extensive aggregations of electron-opaque vesicles were observed near fungal appressoria. These electron-opaque vesicles may be local accumulations of cytoplasm or cytoplasmic aggregates that usually accompany the papilla response (Schmelzer, 2002). These cytoplasmic aggregate-like structures were composed of three types of circular structures similar to those described in Figure 2 (densely staining, lightly staining, and hollow). In addition, we observed numerous circular membrane-bound structures that appeared to be fusing with the plant plasma membrane (red arrows in Fig. 4A1). These were not observed in the compatible interaction. Figure 4B shows the 500^th^ micrograph, where we see another region of local accumulation of plant cytoplasm, which may also be a pre-cursor of papilla deposition (red box). The red arrow again indicates a membrane-bound circular structure appearing to fuse with the plasma membrane. A top view image of the model is shown in Figure 4C, where the extensive interconnected network of papilla-likes structures along the host cell wall is clear. Figure 4D represents a snapshot of the model (Supplementary Video S13) that shows focused deposition of membrane material and reveals a highly interconnected mesh of vesiculo-tubular structures. The appressoria failed to form penetration pegs and no biotrophic hyphae were observed inside host cells. Together these results indicate that when alfalfa is infected with an incompatible *Colletotrichum* species, the fungus fails to penetrate the plant cell, while the plant cell creates an extensive interconnected network of vesiculo-tubular structures underneath attempted fungal penetration sites. Furthermore, large membrane bound vesicles form all along the periphery of the cell that appear to be fusing with the plasma membrane.

**Fig. 4.**
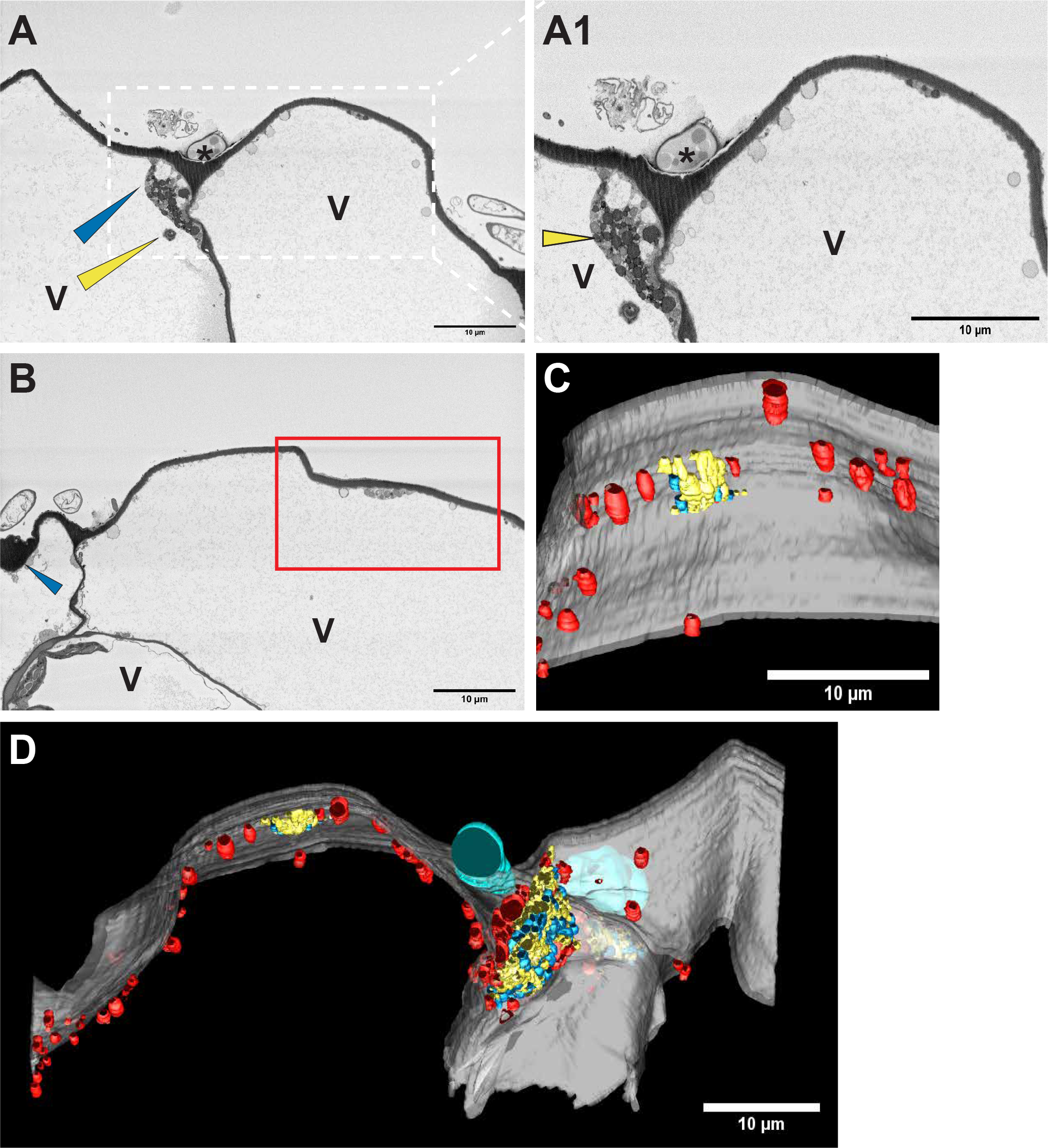
Inoculation with an incompatible pathogen (*C. higginsianum*) induces formation of paramural bodies in alfalfa epidermal cells. **A**, the 1^st^ micrograph, from an SBF-SEM image stack containing 500 images showing alfalfa epidermal cells infected with *C. higginsianum* and imaged 60 hpi. Supplementary Video S12 shows the entire 500 image stack and spans 20 µm of the plant cell. Heavily-stained circular structures are marked with yellow arrowheads and moderately- stained structures are marked with blue arrowheads. The boxed region in panels A is enlarged in panel A1, which shows lightly-stained circles that appear to be fusing to the plant plasma membrane (red arrowheads). **B**, The 500^th^ micrograph from the same image stack. The red arrowhead depicts a circular membrane bound structure that appears to be fusing with the plasma membrane of the host. The red box shows another area of heavily-stained structures similar to those shown in panel A1. **C**, 3D models of the region boxed in red in panel B viewed from top. **D**, 3D model based on all 500 images made using IMOD. Two adjacent plant cell walls are shown in gray. An appressorium is shown in cyan, but no penetration pegs or biotrophic hyphae were observed in any of the 500 images, indicating that the fungus failed to penetrate the host cells. The circular looking vesiculo-tubular structures are modelled in yellow (heavily stained), blue (moderately stained) and red (lightly stained structures that appear to be fusing with the plant plasma membrane). Supplementary Video S13 shows this model rotating in space. V, vacuole; *, appressorium.

To assess whether the cellular changes induced by *C. higginsianum* were associated with a hypersensitive response (HR), we stained cotyledons with trypan blue to detect dead cells. We observed no cell death at either 24 or 60 hpi (Supplementary Fig. S3), indicating that in this interaction resistance is not mediated by HR cell death, possibly because the infection is stopped prior to formation of biotrophic hyphae, hence preventing translocation of effector proteins into host cells. Consistent with the failure to form penetration pegs, we observed increased staining of cell walls with trypan blue (Supplementary Fig. S3), which may reflect increased lignification associated with induction of cell wall-mediated defense responses.

To further assess changes in cell wall structure associated with attempted penetration events, we stained cotyledons with aniline blue, which binds to callose, a polysaccharide that accumulates in papillae that is thought to help block penetration of fungal hyphae (Ellinger et al., 2013; Luna et al., 2011). We observed large fluorescent puncta indicative of callose deposition, which were larger and more abundant in incompatible interactions compared to compatible interactions in both cotyledon and leaves (Supplementary Fig. S4). These results are consistent with the numerous papillae-like structures observed in SBF-SEM images.

### The incompatible fungus *C. higginsianum* induces formation of paramural bodies

The membrane-bound vesicles that appeared to be fusing with the plasma membrane shown in Figure 4 appear similar to previously described paramural bodies (Marchants & Robards, 1968), which form when multivesicular bodies fuse with the plasma membrane. To assess this more carefully, we enlarged a region of the plasma membrane that displayed an abundance of these structures (Fig. 5 and Supplementary Video S14). Red arrows indicate putative paramural bodies. These seem to be a unique feature of the incompatible interaction, as we did not observe any such structures in the compatible interaction or mock-infected sample, suggesting that these structures are associated with immunity. Consistent with this hypothesis, transmission electron micrographs of incompatible interactions of barley and *Arabidopsis* infected with biotrophic powdery mildew fungi have shown multivesicular endosomes and paramural bodies near attempted penetration sites (An, Huckelhoven, et al., 2006), as have TEM images of broad bean plants infected with an incompatible cowpea rust fungus (Xu & Mendgen, 1994). As shown in Figure 4, the paramural bodies observed in the incompatible interaction at 60 hpi are widely distributed along the cell periphery and are not specifically localized at attempted penetrated sites.

**Fig. 5.**
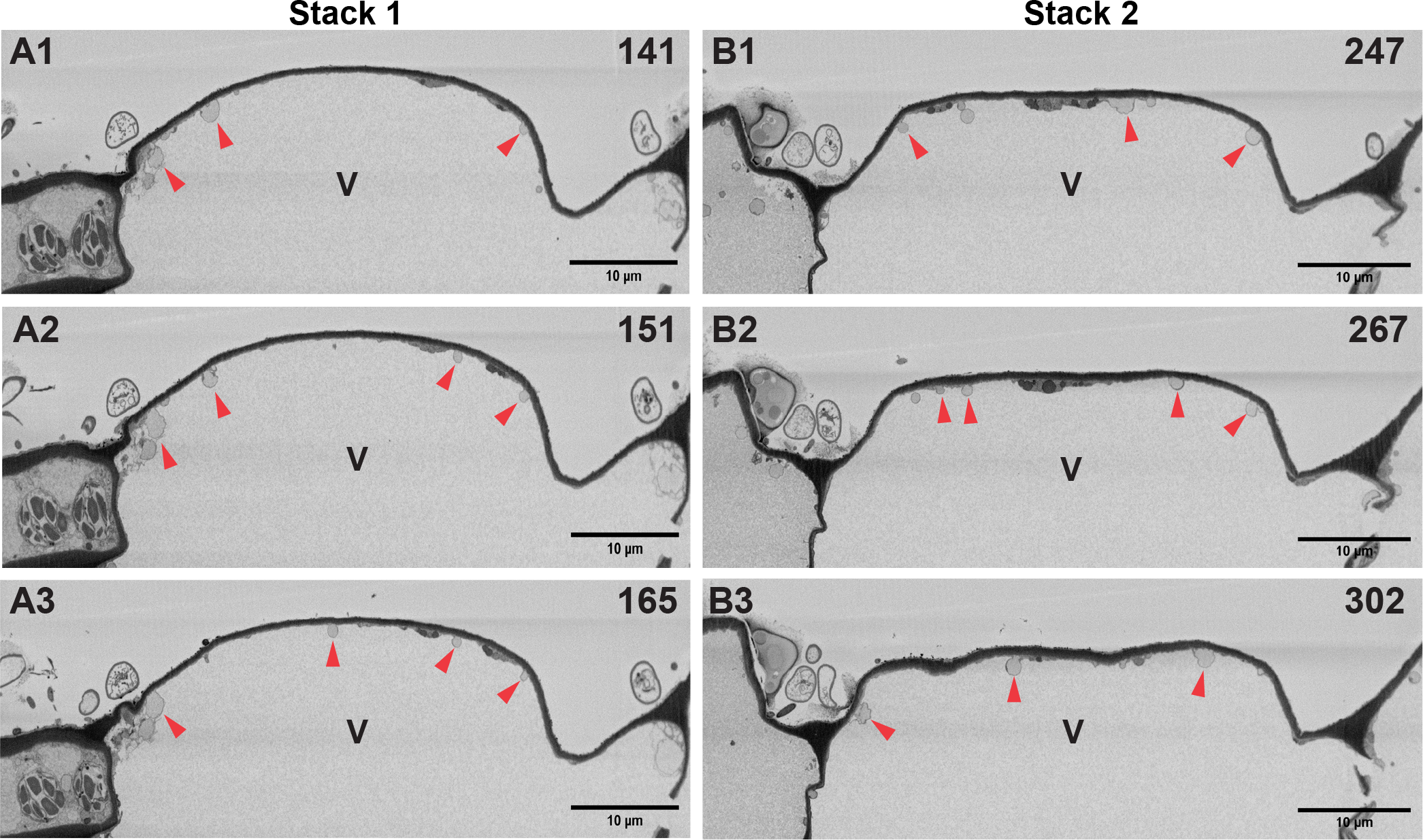
Inoculation with an incompatible fungal pathogen (*C. higginsianum*) induces paramural body-like structures in alfalfa epidermal cells. **A1, A2** and **A3**, The 141^st^, 151^st^ and 165^th^ micrograph from an SBF-SEM image stack containing 500 images showing alfalfa epidermal cells infected with *C. higginsianum* and imaged at 60 hpi (Event 1). **B1, B2** and **B3**, The 247^th^, 267^th^ and 302^nd^ micrograph from the same SBF-SEM image stack showing another event of paramural body fusion to the plasma membrane independent from event 1 (Event 2). Supplementary Video S14 shows all 500 sections. Red arrows indicate paramural body-like structures (circular structures that appear to be fusing with the plasma membrane). V, vacuole.

We quantified the number of paramural bodies observed across the 20 µm of the plant cell imaged by SBF-SEM (Supplementary Fig. S2B). We also calculated the epidermal cell surface area imaged using the SBF-SEM image stack and Fiji software. For block 1 (represented by Supplementary Video S12 and Fig. 4), the surface area was ∼1746 µm^2^ and we counted 174 paramural bodies. For block 2 (represented by Supplementary Video S14 and Fig. 5) the surface area was ∼1478 µm^2^ and we counted 186 paramural bodies. The number of paramural bodies were comparable in the two blocks, although block 2 had more paramural bodies per unit of the cell surface area. To ensure that these PMBs were not an artifact of cell death, we stained infected cotyledons and leaves with trypan blue (Supplementary Fig. S3). These analyses revealed no cell death at 60 hpi for incompatible interactions in either leaves or cotyledons (Supplementary Fig. S3).

Although SBF-SEM imaging enabled us to identify a large number of putative paramural bodies, the xy resolution of SBF-SEM was insufficient to detect structures inside these putative paramural bodies. We therefore used a Focused Ion Beam (FIB)-SEM imaging system, which has an xy resolution of ∼3 nm to better visualize the same resin block shown in Supplementary Figure S1E, Figure 5 and Supplementary Video S14. SBF-SEM uses a microtome to slice thin-sections (40 nm thickness) of the imaging surface parallel to the surface of the stub onto which the resin block is mounted, whereas FIB-SEM uses a gallium ion beam to mill thin layers of the imaging surface perpendicular to the stub surface. The resulting images are thus perpendicular to that produced by the SBF-SEM. Figure 6 and Supplementary Videos S15 and S16 show images obtained using a FIB-SEM. The cell wall, host cell plasma membrane and the tonoplast membrane appear intact. Figure 6C captures the neck region in which an MVB appears to be fusing with the plasma membrane. We used IMOD to reconstruct and model this MVB and adjacent plant membranes (Supplementary Video S17). Figure 6E shows the modelled MVB (blue) above the cell wall (green) with the tonoplast membrane (pink) surrounding the MVB. Figure 6F shows the plasma membrane (white) instead of the tonoplast. Taken together, these images show us that the structures identified by SBF-SEM fusing to the plasma membranes are indeed derived from multivesicular endosomes, as they contain internal vesicles. Supplementary Videos S18 and S19 (enlarged S18) show another MVB with internal vesicles fusing with the PM. These structures are thus paramural bodies and are likely depositing vesicles and other defense-related compounds between the plasma membrane and cell wall.

**Fig. 6.**
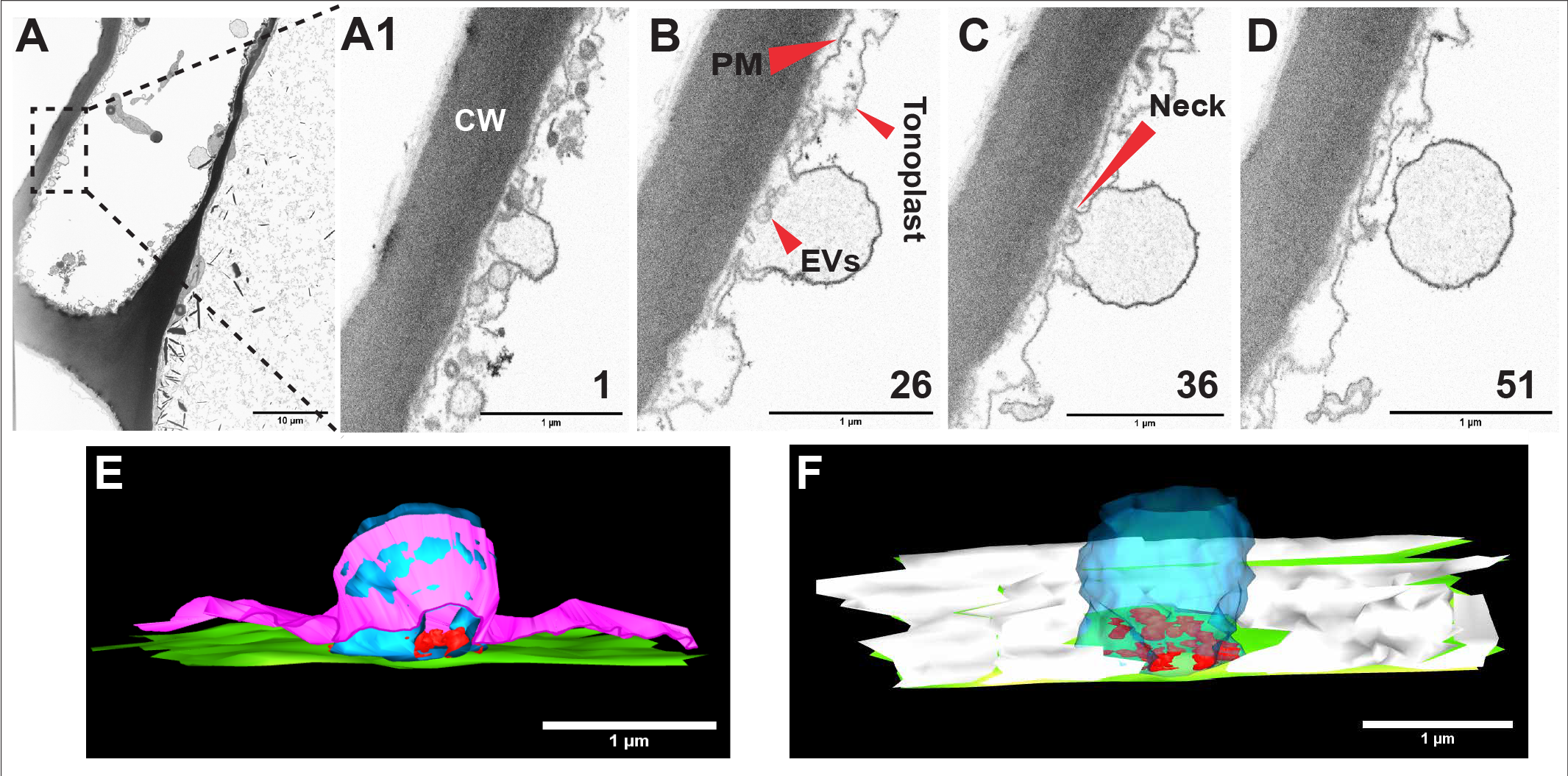
Focused Ion Beam-Scanning Electron Microscope (FIB-SEM) images confirm that *C. higginsianum* induces formation of paramural bodies in alfalfa epidermal cells. **A**, First micrograph from a stack of 51 images obtained using FIB-SEM. Supplementary Video S15 shows the entire stack. Supplementary Video S16 shows the stack with the paramural body region enlarged. The black box shows an area enlarged in panels A1 through D, which highlights a paramural body. **A1**, **B, C**, and **D**, The 1^st^, 26^th^, 36^th^ and the 51^st^ micrographs of the 51-image stack. Supplementary Video S16 shows all 51 images of this region. The plasma membrane (PM), tonoplast, and cell wall (CW) are all labeled. **E**, and **F**, 3D models generated using IMOD of the region shown in panels A1 through D. Green represents the plant cell wall, red highlights the vesicular contents inside the paramural body, blue corresponds to the limiting membrane of the paramural body, pink indicates the tonoplast, and white in panel F and G indicates cell wall. Supplementary Video S17 shows the model of the entire stack rotating in space.

### Fungal infection induces secretion of extracellular vesicles

The numerous paramural bodies induced during the incompatible interaction suggested that large numbers of extracellular vesicles (EVs) should be released. We thus quantified EV release during compatible (*C. destructivum*) and incompatible (*C. higginsianum*) interactions by collecting apoplastic wash fluid and assessing EV content at both 24 hpi and 60 hpi (Fig. 7). We only counted particles between the size ranges of 50-250 nm, which is the typical size range of extracellular vesicles and plotted them in Figures 7A and 7C. At 24 hpi, we did not see a significant increase in vesicle secretion relative to mock infected plants, even though we observed increased numbers of intracellular vesicles at this time point (Figs. 1 and 2).

**Fig. 7.**
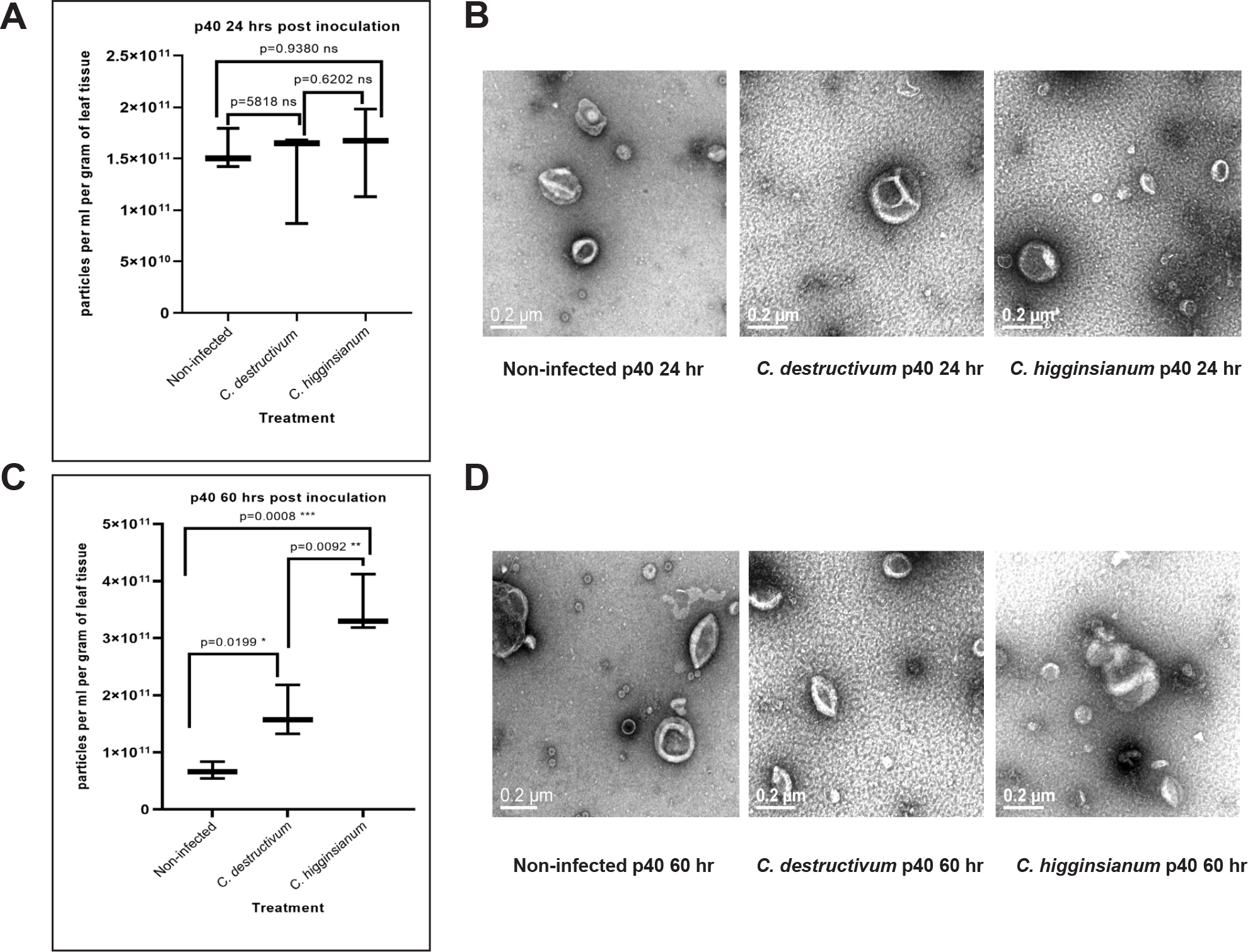
Inoculation with an incompatible fungus induces EV release in alfalfa leaves. **A**, Quantification of extracellular vesicles (EVs) isolated from either mock infected alfalfa leaves, or alfalfa leaves infected with *C. destructium* (compatible) or *C. higginsianum* (incompatible). EVs were isolated by ultracentrifugation of apoplastic wash fluid at 40,000xg (p40) and then quantified using a nanoparticle tracker. The counts of particles between the size range of 50-250 nm are plotted in the graph indicating the true extracellular vesicle population. Data from three biological replicates for each treatment are shown. Error bars indicate standard deviation (SD). p values were calculated using a two-tailed unpaired Student’s T test. **B**, Negative stain TEM images of p40 fractions from representative replicates. **C**, Quantification of EVs (particles between 50-250 nm) isolated at 60 hpi. **D**, Negative stain TEM images of p40 fractions from representative replicates.

By the 60-hour time point, however, we observed a significant increase in EV secretion, with the greatest number of vesicles being secreted in the incompatible interaction, with about a three-fold increase in particle numbers relative to mock infected plants (Fig. 7C). To confirm that we had indeed isolated vesicles, we performed negative stain TEM. Numerous cup-shaped objects (collapsed spheres) ranging in size between 50 and 250 nm diameter were visible, which is typical of EVs. Very few other objects of this size were visible, indicating that the majority of the particles in this size range were EVs. We further confirmed this by performing immunoblots using an antibody to the EV marker protein PEN1 (Rutter & Innes, 2017). Supplementary Figure S5 shows that PEN1 was present in P40 pellets isolated at both 24 and 60 hpi. The increase in EV numbers correlates with the abundance of MVBs and paramural bodies seen in incompatible interactions (Figs. 4A, 4B, Supplementary Fig. S1E, Fig. 5, and Supplementary Videos S12 and S14). Notably, EV numbers observed in the compatible interaction were also higher than mock at 60 hpi (Fig. 7C), suggesting that EVs may also be released via mechanisms that do not involve paramural bodies. Together, these results suggest that EVs are playing a role in the immune response during an incompatible interaction.

## Discussion

We compared the infection process between compatible and incompatible species of *Colletotrichum* on alfalfa by performing serial sections of the plant-fungal interface using SBF-SEM and FIB-SEM. We wanted to answer some key questions: How do these infection structures differ between a compatible and incompatible interaction at an ultrastructural level? Are there any structures unique to the incompatible interaction and vice versa? Can we see the origins of EVs that are released during plant-fungal interactions? The image stacks from these SEM images allowed us to generate 3D models of infection sites, which revealed multiple features that would have been missed using standard TEM. Before detailing the features revealed by 3D modeling, however, we will first discuss the relative strengths and weaknesses of SBF-SEM and FIB-SEM as compared to standard TEM and light microscopy, along with limitations of the present study.

A strength of SBF-SEM relative to both FIB-SEM and standard TEM is the large sample area in three dimensions (individual sample volume) that can be imaged in a relatively short time (e.g., 24 h of microscope time per sample). This enables imaging of multiple cells in a single imaging session in all three dimensions. However, this comes at a cost in terms of resolution. In our case, we were unable to resolve vesicles smaller than about 200 nm using SBF-SEM. We were unable to increase the magnification beyond that shown during our imaging sessions due to charge accumulation on sample blocks induced by the electron beam. Such charge accumulation is particularly problematic with leaf samples due to the large vacuoles in each cell. Vacuoles do not stain well with metal-based EM stains, thus do not conduct electrons well, which results in electron accumulation on the block face at higher magnifications, and hence, poor image quality. To overcome this problem, we employed FIB-SEM, which enables imaging at higher magnifications in all three dimensions compared to SBF-SEM. FIB-SEM also does not suffer from the charge accumulation issue observed with SBF-SEM. However, FIB-SEM is slower than SBF-SEM because each section must be removed by milling rather than with a microtome, and the total area that can be imaged in single image is much less. The increase in required microscope time typically means an increase in expense.

Both SBF-SEM and FIB-SEM greatly facilitate 3D reconstructions compared to standard TEM because there is no need to collect and image serial sections that have been sliced off of a block, which can be very challenging, especially when collecting hundreds of sections as was done in the present study. Because each image from SBF-SEM and FIB-SEM is generated from a block face, it is also easier to align serial images, compared to images of independent sections generated for standard TEM, which often have artifacts induced by uneven shrinkage or expansion of ultrathin sections. That said, TEM still offers the highest possible resolution, and may be necessary when one needs to image structures smaller than 10 nm.

Although the above EM technologies offer outstanding resolution compared to light microscopy, especially of live cells, the present study is limited by the fact that we are imaging fixed, resin-embedded cells and are thus using static images to understand a dynamic process. This is particularly relevant to assessing whether vesicles that are fused to the plasma membrane are in the process of endocytosis or exocytosis. In this study, we have assumed that all such vesicles observed by SBF-SEM are in the process of exocytosis. This assumption is supported by several observations. First, higher resolution images obtained using FIB-SEM revealed the presence of extracellular vesicles outside the plasma membrane immediately adjacent to these fusions. It is unlikely that such extracellular vesicles would be present during endocytosis. Second, these presumptive paramural bodies are quite large, pushing into the vacuole against the turgor pressure of the vacuole. It is unclear how endocytosis could generate such a large invagination of the plasma membrane without anything pushing from the outside. Lastly, these paramural bodies were specifically induced by inoculation with an incompatible fungus, which we showed resulted in a large increase in vesicle secretion. It seems unlikely that, at the same time, the cells would be undergoing a large increase in a highly unusual form of endocytosis.

Although SBF-SEM is limited in resolution compared to standard TEM, we were still able to resolve extensive accumulation of circular structures along the periphery of inoculated cells induced by both compatible and incompatible *Colletotrichum* species. Most of the circular structures were electron-opaque, while some were lightly stained. In addition, the incompatible interaction also accumulated circular structures with no internal staining, which were not present in the compatible interaction at this time point. By using SBF-SEM, we were able to trace these circular structures through the volume of the plant cell to see if they were vesicular or tubular in 3D. This work revealed that many were, in fact, tubular and were interconnected, forming a mesh-like network. Previous studies that employed standard TEM to image fungal infection sites on plant cells have revealed similar circular structures, which were assumed to be spherical vesicles or organelles (O’Connell, 1987; O’Connell et al., 1985; Politis, 1976; Wharton et al., 2001), but from our SBF-SEM observations we now expect these may have been cross-sections through more complex structures. Regardless, accumulation of vesicles and tubules along the periphery of inoculated cells appears to be an immune response, as they were not seen in non-infected control leaves.

By 24 hpi, both compatible and incompatible *Colletorichum* species had formed appressoria while neither fungal species had penetrated the host cell wall. This is consistent with previous reports of compatible interactions between cucumber leaves and *Colletotrichum lagenarium* (Xuei et al., 1988), maize leaves and *C. graminicola* (Mims & Vaillancourt, 2002) and bean leaves and *C. lindemuthianum* (O’Connell et al., 1985).

At 60 hpi with the compatible pathogen, we observed penetration pegs beneath appressoria that gave rise to bulbous biotrophic hyphae. These hyphae displaced the majority of the plant cell central vacuole. 3D modelling enabled us to see the large volume occupied by these hyphae and their branching. This would not be posible with standard TEM. This finding is consistent with TEM-based observations of *C. lindemuthianum* infecting epidermal cells of French bean, which showed cross-sections of large biotrophic hyphae displacing epidermal cell central vacuoles at 4 days post inoculation (O’Connell et al., 1985). Similar bulbous biotrophic hyphae that arise from appressoria have also been observed with *C. sublineolum* infecting sorghum epidermal cells (Wharton et al., 2001), and in multiple other compatible *Colletotrichum*- plant interactions using both light and electron microscopy. For example, the *C. gloeosporioides* – tangerine interaction (Brown, 1977), *C. graminicola* – maize interaction (Mims & Vaillancourt, 2002), and *C. lagenarium* (now called *Colletotrichum orbiculare*) – cucumber interaction (Xuei et al., 1988).

In contrast to the compatible interaction, we did not observe any penetration pegs or biotrophic hyphae in the incompatible interaction at 60 hpi. This observation is similar to that observed in *Arabidopsis* infected with incompatible (non-adapted) *Colletotrichum* species (Shimada et al., 2006), which indicates that incompatible fungi are typically stopped prior to penetration of the host cell wall without induction of host cell death. This differs from resistant interactions to compatible (adapted) species of fungi, which typically are induced following penetration of the host cell and secretion of effectors (Howlett et al., 2012; Irieda et al., 2014). For example, small intracellular hyphae with no interfacial matrix were observed in resistant bean varieties infected with *C. lindemuthianum* (O’Connell et al., 1985). Similarly, formation of penetration pegs with occasional biotrophic hyphae have been observed in resistant sorghum infected with *C. sublineolum* (Wharton et al., 2001), resistant oats infected with *C. graminicola* (Politis, 1976), and resistant cucumber infected with *C. lagenarium* (Xuei et al., 1988).

Although we did not observe host cell death or penetration of the host cell wall during incompatible interactions, extensive production of electron-opaque vesicle-like structures was seen, mostly beneath appressoria. 3D modelling revealed these structures to be compact layers of interconnected membranes. This meshwork is likely part of a local accumulation of host cytoplasm associated with papilla formaton. Papilla formation occurs in response to several *Colletotrichum* species during both host and non-host interactions. For example, papilla formation was observed in resistant and susceptible *Medicago sativa* infected with *C. trifolii* (Mould et al., 1991; Mould & Robb, 1992), cucumber plants infected with virulent or avirulent *C. lagenarium* strains (Xuei et al., 1988), sorghum infected with virulent or avirulent *C. sublineolum* (Wharton et al., 2001), French bean infected with virulent or avirulent *C. lindemuthianum* (O’Connell et al., 1985), *Arabidopsis* infected with a non-host *Colletotrichum* species (Shimada et al., 2006), and oats infected with avirulent *C. graminicola* (Politis, 1976). Although papillae fail to stop infection of virulent Colletotrichum strains, it is thought that they contribute to resistance against avirulent and non-host strains. This is based, in part, on differences in the appearance of papillae formed during resistant and non-host interactions versus those formed during susceptible interactions. For example, resistant sorghum produces papillae that are more electron-opaque compared to those formed by a susceptible variety, and papillae in the resistant variety are penetrated at a lower rate compared to those of the susceptible variety (Wharton et al., 2001). Similarly, penetration of papillae formed in cucumber occurs at a much lower frequency during infection by avirulent strains than during infection by virulent strains (Xuei et al., 1988).

In addition to revealing the complex membrane arrangements present in the papilla precursors or cytoplasmic aggregates, our 3D EM analyses revealed a dramatic increase in plant paramural body formation during the incompatible interaction with *C. higginsianum* compared to the compatible interaction with *C. destructivum*. To our knowledge, this is the first report of plant paramural bodies accumulating in response to challenge by a non-adapted or incompatible fungal pathogen, and the finding suggests that paramural bodies could contribute to the *Medicago* defense response.

Paramural bodies are thought to be formed by the fusion of multivesicular bodies (MVBs) with the plasma membrane, thereby depositing the vesicles, and any other components found inside the limiting membrane of the MVB, outside the plasma membrane. Paramural bodies have been observed previously near the vicinity of papillae during both compatible and incompatible interactions with both hemibiotrophic *Colletotrichum* as well as with the biotrophic powdery mildew fungus *Blumeria graminis* f. sp. *Hordei* (An, Ehlers, et al., 2006; An, Huckelhoven, et al., 2006). Our SBF-SEM data revealed, however, that paramural bodies were broadly distributed around the periphery of the cell, and were not limited to the region around papillae.

The increase in paramural body formation during the incompatible interaction suggested that there should be an increase in extracellular vesicle release. We therefore isolated extracellular vesicles from both compatible and incompatible interactions at both 24 hour and 60 hour time points. These analyses revealed that, compared to mock infected plants, EV abundance increased more than three-fold by 60 hpi with the incompatible strain and approximately two-fold with the compatible strain. The increase observed with the compatible strain in the absence of an obvious increase in paramural bodies suggests that plants may also release EVs using a mechanism independent of paramural bodies. The increase in EV abundance with both compatible and incompatible strains is consistent with our previous findings that plants sprayed with salicylic acid or infected with *Pseudomonas syringae* increase EV abundance approximately two-fold (Rutter & Innes, 2017).

Taken together, our results suggest that paramural bodies and papillae contribute to non-host resistance in plants and that the contents of paramural bodies, including EVs, may play a role in such resistance. Further study is required to investigate the contents of the EVs released during incompatible interactions and how these differ from EVs isolated from uninfected plants.

## Materials and Methods

### Fungal material

*Colletotrichum destructivum* isolate CBS 520.97 (https://wi.knaw.nl/Collection) and *Colletotrichum higginsianum* isolate IMI349063A were used for infections. The fungal stocks were stored at -80°C in 1X potato dextrose broth supplemented with 20% glycerol. Fungal cultures were prepared by spotting cultures from glycerol stocks onto solid Mathur’s Media (2.8 g/L glucose, 2.2 g/L mycological peptone, 0.5 g/L Yeast Extract, 1.2 g/L MgSO4x7H2O, 2.7 g/L KH2PO4, 20 g/L agar, pH 5.5). Plates were kept in the dark for 24 hours (h) and then allowed to grow for 3 weeks under short day photoperiod conditions (9h days, 22°C, and 150 µEm^−2^s^−1^). For harvesting the spores, 1 mL of sterile deionized water was applied to the Mathur’s plate, and the spores were dislodged using a sterilized glass rod. The spore solution was transferred to a microfuge tube and the spores were then pelleted at 2000xg for 5 minutes (min), discarding the supernatant containing the mycelia. The spore pellet was washed five times with 1 mL deionized water, centrifuging each time at 2000xg for 5 min. The final pellet was resuspended in sterile deionized water and spores were counted using a Neubauer chamber and adjusted to the desired concentration.

### Plant material and growth conditions for spot inoculation and vesicle isolation

Alfalfa (*Medicago sativa*) seeds were sterilized using 10% bleach (30 seconds) and 70% ethanol (2 min) followed by five rinses in sterile deionized water. The sterilized seeds were placed on Pro-mix PGX potting mix inside a humidity chamber that was created by placing pots inside a plastic box which in turn was placed inside a plastic bag. The seeds were stratified at 4°C for 24 h before moving them to a growth room (10h days, 24°C, and 150 µEm^−2^s^−1^) for 7 days. Each cotyledon was then spot inoculated with ∼5 µL of 2x10^6^ spores/mL (mixed with 0.001% Silwet) of either *Colletotrichum higginsianum* (incompatible) spores or *C. destructivum* (compatible) spores. Inoculated plants were incubated in the dark for 12 h, at 100% humidity at 25°C before being transferred to the short-day room. For mock-inoculated controls, ∼5 µL of sterile deionized water (mixed with 0.001% Silwet) was spotted onto each cotyledon under identical conditions as the infected samples. The water droplet or the fungal droplet did not evaporate even after 60 hours, suggesting that the humidity chamber maintained 100% humidity.

For vesicle isolation, alfalfa seeds were sterilized and stratified as described above and then grown in a growth room as described above for 3 weeks before being sprayed with either *C. higginsianum* or *C. destructivum* spores (2x10^6^ spores/mL mixed with 0.001% Silwet), or water (mixed with 0.001% Silwet) as a control. The sprayed plants were covered with a plastic dome and then placed inside plastic bags to maintain humid conditions until vesicle isolation at either 24 hpi or 60 hpi.

### Sample fixation, staining and resin embedding

Inoculated regions of alfalfa cotyledons were sampled using a 2 mm biopsy punch at either 24 hpi or 60 hpi. Several 2 mm discs were collected for each infection condition and the discs were then placed in an 8 mL glass sample vial with rubber lined cap (Wheaton, Catalog # 224884) and vacuum infiltrated with fixative solution composed of 4% glutaraldehyde (v/v, EM Grade, Electron Microscopy Sciences, Cat# 16020) and 4% formaldehyde (v/v, EM Grade, EMS, Cat# 15710) in 100 mM sodium phosphate (SP) buffer pH 7.2 until the samples sank to the bottom. Infiltrated samples were incubated at 4°C under constant rotation for 24-36 hours. The sample was then washed five times for 10 min each with SP buffer and transferred to a freshly prepared solution containing 1.5% potassium ferrocyanide (w/v, SIGMA Life Science, Lot# BCBV7953) and 2% OsO4 (v/v, EM Grade, EMS, Cat# 19150) solution in SP buffer for 2 h at room temperature while rotating. Subsequently, the samples were washed five times for 10 min each with sterile deionized water and then transferred to fresh solution containing 1% (w/v) thiocarbohydrazide (EMS, Cat# 21900) in deionized water and incubated for 1 hour under rotation at room temperature. The samples were then washed with deionized water five times for 10 min and incubated with 2% OsO4 (v/v) in deionized water for 2 hours at room temperature. The samples were again washed with deionized water five times for 10 min before staining with 0.2% (w/v) uranyl acetate (EMS, Cat# 22400) in water overnight at 4°C. The samples were again washed five times with deionized water for 10 min each, before gradually dehydrating with 10% (v/v), 25%, 50%, 75%, and 3 x 100% acetone for 30 min each. Samples were then infiltrated with Durcupan resin (EMS, Cat# 14040) in 10% (v/v in acetone), 25%, 50%, 75%, and 5 x 100% resin over a period of 3 days. Samples were carefully layered on top of fresh resin during each of the 100% resin infiltration steps and centrifuged at 376xg until the samples reached the bottom of the 2 mL centrifuge tubes (usually ∼2 min) (McDonald, 2014). Samples were finally flat-embedded in Durcupan using Aclar® (Kingsley & Cole, 1988) or flat-embedding molds in an oven at 60°C.

### Sample preparation for electron microscopy

Samples embedded in resin were first trimmed roughly using a jeweler’s saw and then mounted on aluminum stubs using EPO-TEK® electrically conductive resin, which was allowed to solidify in an oven at 60°C overnight. Samples were then trimmed by hand using a double-edged razor blade to an approximate height of 0.5 mm, length of 2 mm, and width of 300 µm, followed by sputter-coating with 45 nm 80:20 Au:Pd at 3.8x10^-2^ Torr and 30 mA using Safematic CCU-010 Compact Coating Unit, before smoothening the block-face with a diamond knife (Diatome) using a Leica Ultracut ultramicrotome.

### Serial block-face scanning electron microscopy

Samples were imaged using a Thermo-Fisher Teneo Volume Scope equipped with an in situ ultramicrotome, which was set to remove 40 µm sections with each slice. Images of the block face were obtained at 2.5 kV/0.8 nA at a low vacuum pressure of 50 pa with the VS-DBS detector for backscatter. Each of the images were 6144 x 4096 pixels with an x-y pixel size of 10 nm. The images obtained were first batch inverted using Adobe Photoshop CC, and then processed using IMOD (Kremer et al., 1996). This software was used to perform segmentation and 3D reconstructions. Videos were created from serial images using the Windows 10 Media player function.

### Focused ion beam-scanning electron microscopy

FIB-SEM image acquisition was performed using a Zeiss Auriga 60 FIB-SEM with ATLAS 3D software (FIBICs, Carl Zeiss Microscopy). We used a sample preparation workflow containing the following steps: (I) depositing a 30 × 20 µm platinum pad of 1 µm thickness on the sample surface over the region of interest using the gas injection system (GIS); (II) milling tracking and autotune marks in the platinum pad using the Ga ion beam; (III) depositing a carbon layer of 1 µm thickness onto the tracking and autotune marks for protection during the milling and imaging process; (IV) coarse trench milling (30 µm width and 30 µm depth); and (V) trench polishing (30 µm width and 30 µm depth). The SEM images were acquired at 1.5 kV with the energy-selective back-scattered electron (EsB) detector at a grid voltage of 750 V and 1 nA of electron beam current (60 µm aperture and high current mode). The pixel dwell time was 1 µs with line average of 1. The x-y pixel size was 3 nm and the slicing thickness in the z-direction was 12 nm. During the milling and imaging process, the FIB ion beam was operated at continuous milling mode at 30 kV and 600 pA of ion beam current with a milling rate of 8 nm/min. About 700 images were acquired continuously in a period of about 19 hours.

### Vesicle isolation

Apoplastic wash fluid was isolated from 3-week-old alfalfa plants following the protocol described in (Rutter & Innes, 2017). In brief, leaves were vacuum infiltrated with Vesicle Isolation Buffer (VIB, 20 mM MES, 2 mM CaCl2, and 0.1 M NaCl, pH6). The excess buffer was removed from the surface of the leaves using Kimwipes®. The leaves were then placed inside needleless 30 mL syringes (making sure not to overstuff the syringes) and the syringes were placed inside 50 mL screwcap tubes. The tubes were then centrifuged for 20 min at 700 x *g* with slow acceleration at 4°C using a JA-14 rotor (Avanti J-20 XP centrifuge; Beckman Coulter). The resulting apoplastic wash collected was passed through a 0.22 µm membrane filter to remove bacteria and larger debris. The supernatant was transferred to new 13 x 51 mm polycarbonate centrifuge tubes (Beckman Coulter # 349622) and centrifuged at 10,000 x *g* for 30 min at 4°C using a TLA100.3 fixed angle rotor and an Optima TLX Ultracentrifuge (Beckman Coulter).

Supernatants were then transferred to new centrifuge tubes and centrifuged at 40,000 x *g* for 1 hour at 4°C using the same rotor. The resulting pellet was washed using VIB and re-pelleted at 40,000 x *g* for 1 hour at 4°C. The final pellet was resuspended in 50 µL of fresh filtered VIB and kept on ice for further experiments or stored at -80°C for subsequent use.

### NanoParticle tracking analysis

Particle concentrations and diameters were determined using a ZetaView Particle Tracking Analyzer (ParticleMetrix, Diessen) and its built-in software. The analyzer was first calibrated using 100 nm polystyrene beads. Vesicle samples were diluted 1:1000 times using fresh VIB pH 6.0. The Zetaview was operated with a max diameter setting of 500 nm, and minimum diameter setting of 5 nm. For data analysis, the percentage of particles that were between 50-250 nm in diameter was calculated and plotted in the graph.

### Transmission electron microscopy of extracellular vesicles

Aliquots (5 µL) of resuspended P40 pellets were spotted onto Formvar and carbon-coated copper electron microscopy grids (Electron Microscopy Sciences) that were glow-discharged at 15mA for 60 seconds prior to application of samples. After 5 min, the excess sample was wicked off using filter paper. Following that, 10 µL of 2% uranyl acetate was applied to the grids for 2 min. The excess solution was wicked off using filter paper and the grids were allowed to air dry overnight. The grids were imaged using a JEM-1010 transmission electron microscope (JEOL USA) at 80 kV.

### Trypan Blue Staining

Alfalfa cotyledons and leaves were first stripped of their chlorophyll by incubating in 1:3 w/v solution of acetic acid:95% ethanol overnight with gentle shaking. For staining, a stock solution of trypan blue (10 g phenol, 10 ml lactic acid, 10 ml glycerol, 10 ml de-ionized water, and 0.02 g of trypan blue (Sigma-Aldrich- 302643-25G)) was diluted with 95% ethanol (1:2 v/v). Samples were placed in a 55°C water bath with 1 ml of trypan blue solution for 1 minute and then transferred to 4°C overnight with gentle shaking. To destain, chloral hydrate solution was prepared by mixing 1000 g of chloral hydrate (Sigma-Aldrich- 302-17-0) in 400 ml de-ionized water (takes several hours to dissolve). Samples were incubated in chloral hydrate solution for a minimum of 3 h with replacement of the solution approximately once per hour.

### Aniline Blue Staining

Alfalfa cotyledons and leaves were first stripped of their chlorophyll by placing them in 1:3 w/v solution of acetic acid:95% ethanol overnight with gentle shaking. The samples were then subsequently rehydrated with 150 mM phosphate buffer, pH 8.0 for 30 minutes. Samples were stained in 150 mM phosphate buffer pH 8.0 containing 0.01% (w/v) aniline blue (Sigma Aldrich- 28631-66-5) at 4°C overnight. Samples were imaged using a Nikon NiE microscope equipped with a DAPI filter.

### Immunoblots

For immunoblot analysis, P40 pellets were resuspended in 50 µl of VIB buffer and 20 µl of this suspension was mixed with 5 µL of 5X SDS loading buffer (8% SDS, 250 mM Tris-HCl, pH 6.8, 0.1% Bromophenol Blue, 40% glycerol, and 400 mM dithiothreitol) and heated at 95°C for 5 min. Cell lysates (positive controls) were prepared by freezing 500 mg of leaf tissue in liquid nitrogen and grinding with a mortar and pestle. Ground leaf tissue was extracted in 1 mL of protein extraction buffer (50 mM Tris-HCl pH 7.0, 150 mM NaCl, 0.1% Nonidet P-40, 1% plant protease inhibitor cocktail (Sigma-Aldrich) and 1% 2,2’-dipyridyldisulfide) and centrifuged at 12,500 x *g* for 10 minutes at 4°C to pellet debris. 20 µL of supernatant was then mixed with 5 µL of 5X SDS loading buffer and heated at 95°C for 5 min.

Samples were loaded on 4% to 20% Precise Protein Gels (ThermoScientific) and electrophoresed at 120V for 1 h in Tris-SDS running buffer (30.0 g of Tris base, 144.0 g of glycine, and 10.0 g of SDS in 1000 ml of H2O, pH 8.3 (10x) diluted to 1x before use). The proteins were transferred to a nitrocellulose membrane (Amersham Protran Premium 0.45 µm NC product 10600003). Ponceau staining was used to confirm successful transfer and equal loading of samples. Membranes were washed with Tris-buffered saline (50 mM Tris-Cl and 150 mM NaCl, pH 7.5) containing 0.1% Tween 20 (TBST) and blocked with 5% Difco Skim Milk (BD) overnight at 4°C. Membranes were incubated with anti-PEN1 antibody (Zhang et al., 2007) at a 1:1,000 dilution for 1 h, washed with TBST, and incubated with horseradish peroxidase-labeled goat anti-rabbit antibody (Abcam AB97051) at a 1:5,000 dilution for 1 h. After a final wash in TBST, protein bands were imaged using a BIO-RAD ChemiDoc Imaging system.

## Supplementary Videos

All supplementary videos can be viewed at the following link: https://drive.google.com/drive/folders/1AGh3bU59uwFenLad0YESg44gZMSav6wA?usp=sharing

## ACKNOWLEDGEMENTS

We thank the IU Bloomington Electron Microscopy Center at Indiana University for access to A Leica Ultracut ultramicrotome, JEOL JEM 1010 Transmission Electron Microscope, and ThermoFisher Teneo VolumeScope. The latter microscope was funded by an equipment grant from the National Institutes of Health (grant number 1S10OD023501). We also thank the IU Bloomington Nanocharacterization Facility (NCF) for access to a Zeiss Auriga 60 FIB-SEM and to NCF/IUB-EMC staff member Jun Chen for assistance with this equipment.

---

**Supplementary Fig. S1.**
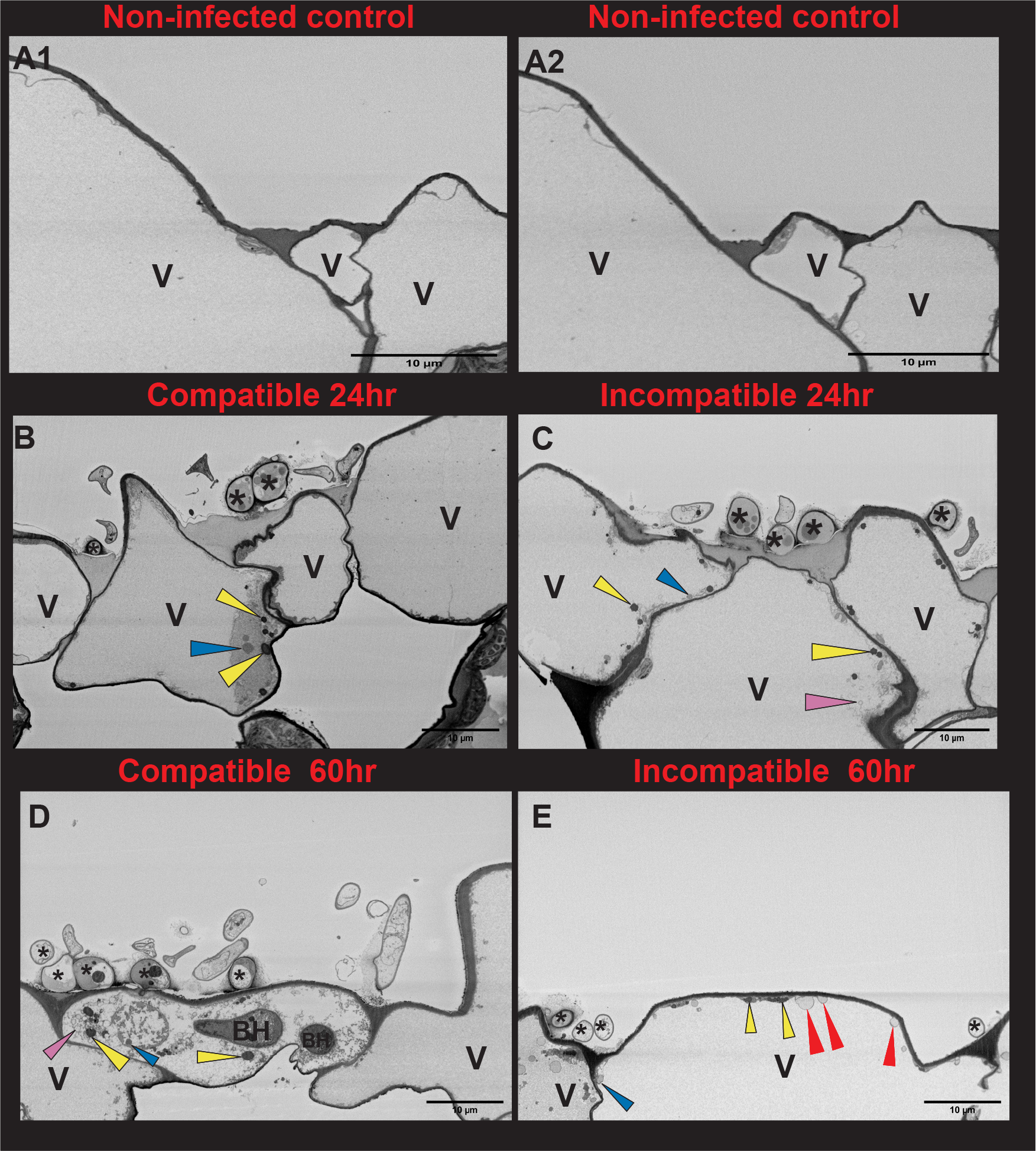
Inoculation with an incompatible fungal pathogen (*C. higginsianum*) induces paramural bodies in alfalfa epidermal cells. **A1** and **A2**, The 1^st^ and the 101^st^ micrograph from an SBF-SEM image stack containing 200 images showing alfalfa epidermal cells mock infected with a water droplet and imaged 60 hour after inoculation. Supplementary Video S1 shows all 200 sections. **B**, 100^th^ micrograph from an SBF-SEM image stack containing 400 images showing alfalfa epidermal cells infected with *C. destructivum* (a compatible species) and imaged 24 hours after inoculation. Supplementary Video S11 shows all 400 sections. **C**, 60^th^ micrograph from an SBF-SEM image stack containing 400 images showing alfalfa epidermal cells infected with *C. higginsianum* (an incompatible species) and imaged 24 hours after infection. Supplementary Video S4 shows all 400 sections. **D**, 1^st^ micrograph from an SBF-SEM image stack containing 400 images showing *alfalfa* epidermal cells infected with *C. destructivum* and imaged 60 hours after inoculation. Supplementary Video S11 shows all 400 sections. **E**, 237^th^ micrograph from an SBF-SEM image stack containing 500 images showing alfalfa epidermal cells infected with *C. higginsianum* and imaged 60 hours after inoculation. Supplementary Video S12 shows all 500 sections. Heavily-stained circular structures are marked with yellow arrowheads. Moderately-stained circular structures are marked with blue arrowheads and lightly-stained circular structures marked with pink arrowheads. Red arrowheads mark circular structures that appear to be fusing with the plasma membrane. BH, biotrophic hyphae; V, vacuole; *, appressoria.

**Supplementary Fig. S2.**
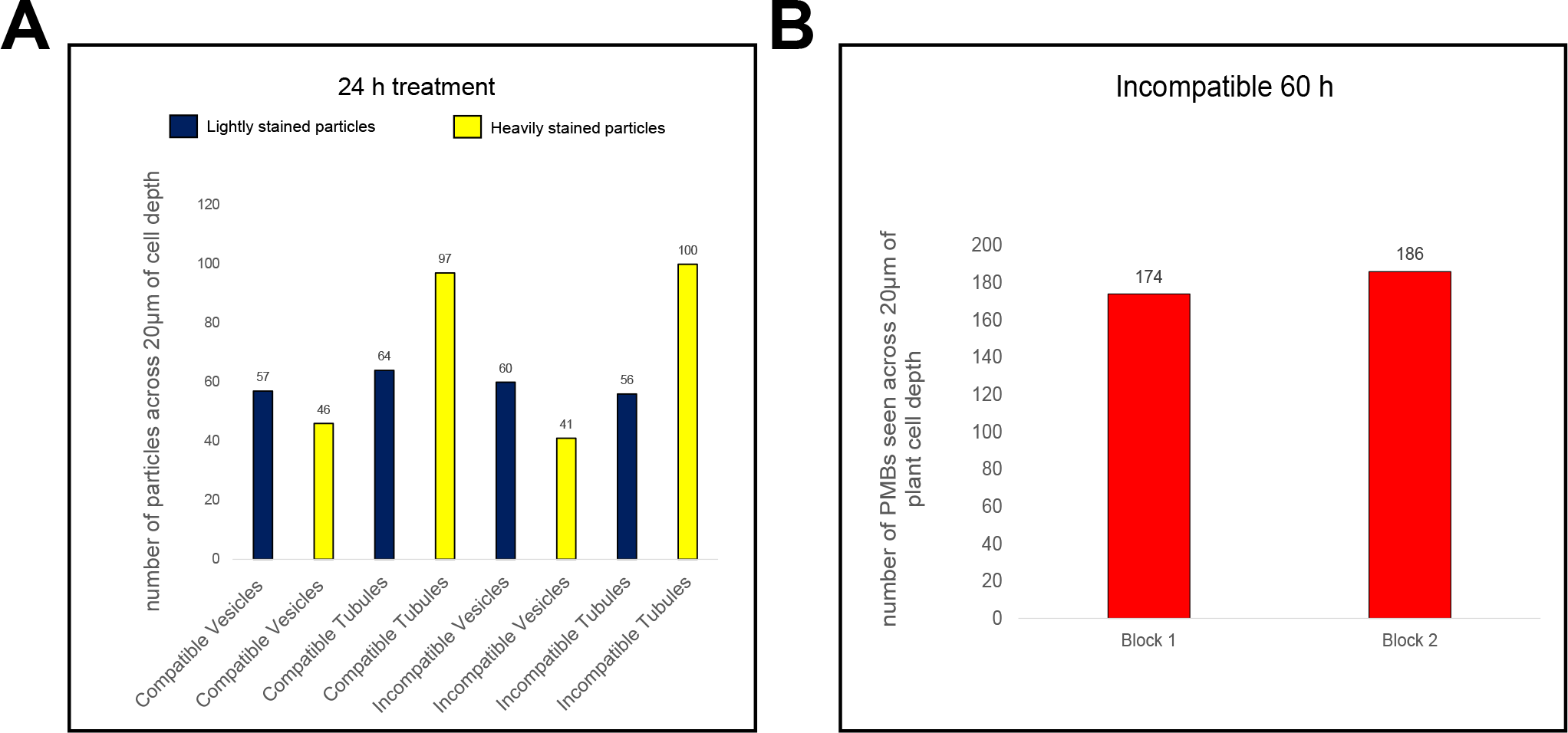
Quantification of vesicles and tubules observed in compatible and incompatible interactions at 24h, and paramural bodies observed in incompatible interactions at 60h. **A**, Graph showing the number of particles (vesicles or tubules) observed across 20 µm of plant cell depth in alfalfa cotyledons infected with *C. destructivum* (compatible) or *C. higginsianum* (incompatible) 24 h post infection. Blue represents moderately stained particles and yellow represents heavily stained particles. **B**, Graph showing the number of paramural bodies observed across 20 µm of plant cell depth at 60 h post infection with incompatible *C. higginsianum*. The two bars represent observations for two separate blocks. Supplementary Video S12 represents block 1 and Supplementary Video S14 represents block 2.

**Supplementary Fig. S3.**
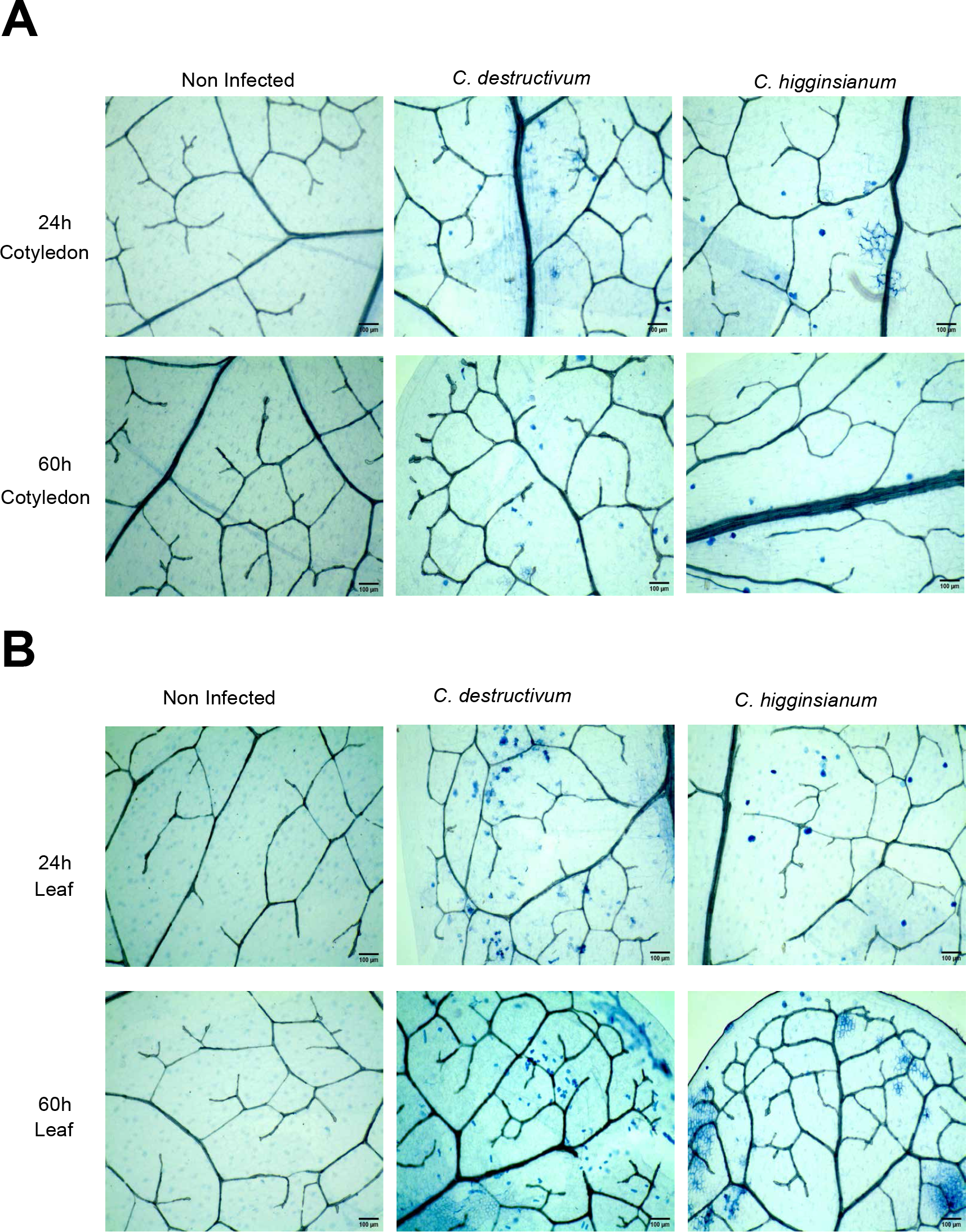
Alfalfa cotyledons and leaves infected with either compatible (*C. destructivum)* or incompatible (*C. higginsianum*) *Colletotrichum* species show little to no cell death at either 24 h or 60 h post inoculation. **A**, Alfalfa cotyledons were infected with the indicated fungal strains and then stained with trypan blue at 24 h or 60 h post inoculation. Vascular cells are stained, but no staining of mesophyll or epidermal cells is observed. **B**, Alfalfa leaves were infected with the indicated fungal strains and then stained with trypan blue at 24 h or 60 h post inoculation. Vascular cells are stained, as are biotrophic hyphae of *C. destructivum* at 60 h, but no staining of mesophyll or epidermal cells is observed. Some staining of anticlinal cell walls is observed in samples infected with *C. higginsianum*.

**Supplementary Fig. S4.**
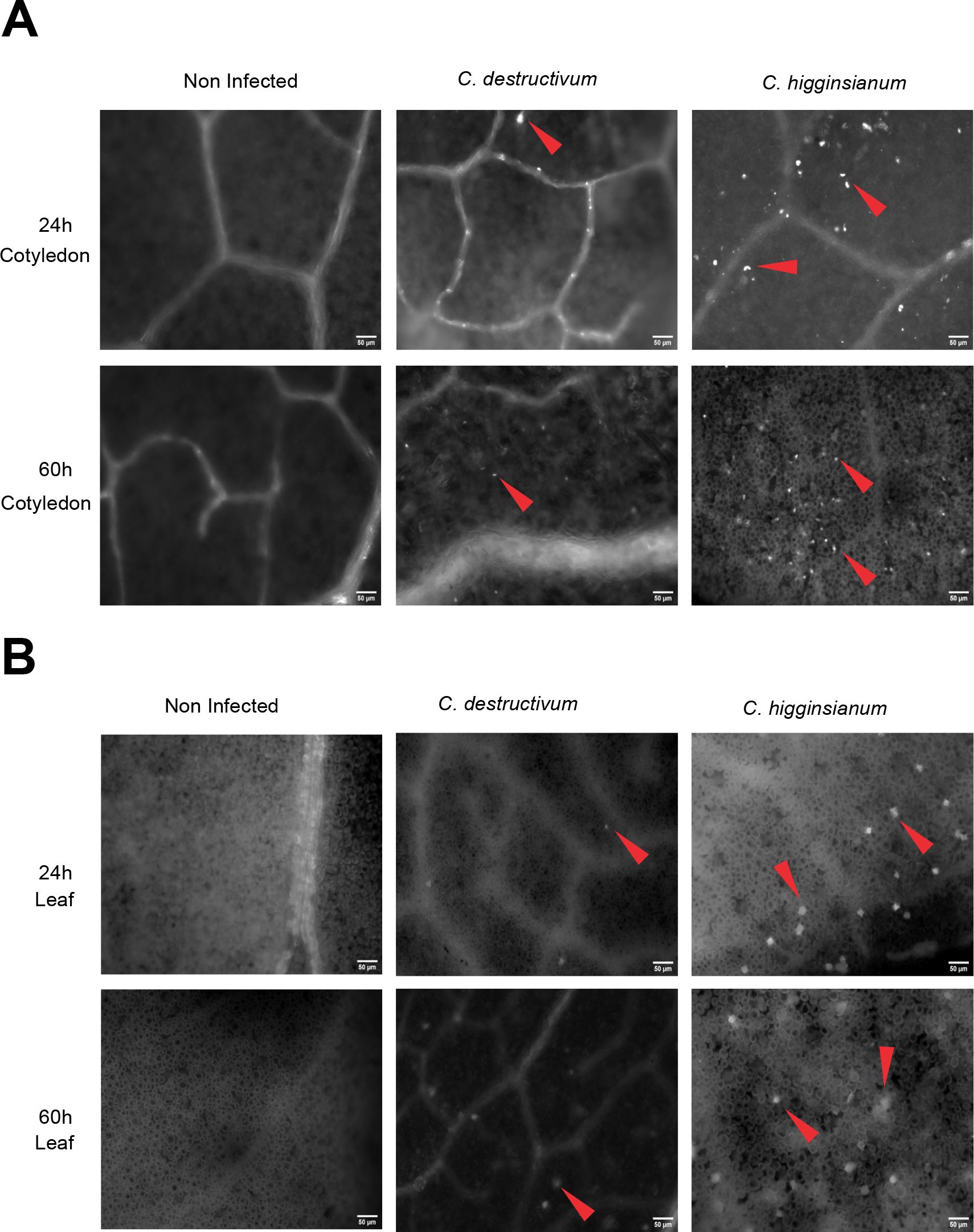
*C. higginsianum* induces callose deposits in alfalfa, a non-host. **A**, Alfalfa cotyledons were infected with the indicated fungal strains and then stained with aniline blue at 24 h or 60 h post inoculation. Very few callose deposits were observed in the non-infected control or in cotyledons infected with the compatible strain, *C. destructivum*, whereas many deposits were observed in cotyledons infected with the incompatible strain, *C. higginsianum*. **B**, Alfalfa leaves were infected with the indicated fungal strains and then stained with aniline blue at 24 h or 60 h post inoculation. Very few callose deposits were observed in the non-infected control or in leaves infected with the compatible strain, *C. destructivum*, whereas many deposits were observed in leaves infected with the incompatible strain, *C. higginsianum*.

**Supplementary Fig. S5.**
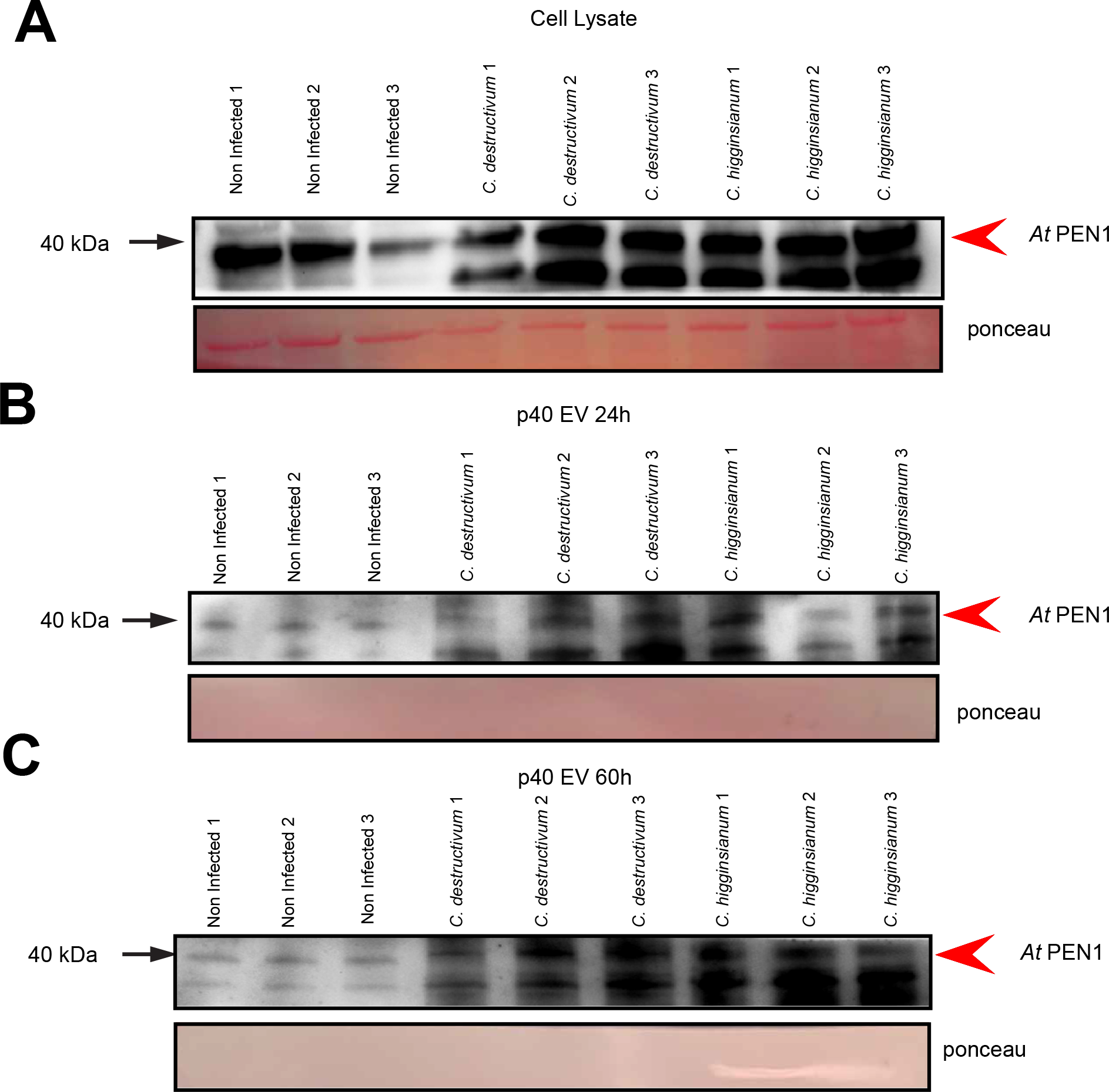
P40 pellets isolated from alfalfa leaves contain the EV marker protein PEN1. **A**, Anti- PEN1 immunoblot of cell lysate isolated from three replicates of non-infected, compatible (*C. destructivum)* or incompatible (*C. higginsianum*) infected plants. Bottom panel shows Ponceau-stained blot to assess equal loading. **B**, Anti-PEN1 immunoblot of P40 pellets isolated 24 h post inoculation with compatible (*C. destructivum)* or incompatible (*C. higginsianum*) *Colletotrichum* species. Bottom panel shows ponceau stained blot. Lack of a Rubisco band indicates P40 pellet has very little contamination with non-vesicle proteins. **C** Anti- PEN1 immunoblot of P40 pellets isolated 60 h post inoculation.

